# Parallel systems for social and spatial reasoning within the cortical apex

**DOI:** 10.1101/2021.09.23.461550

**Authors:** Ben Deen, Winrich A. Freiwald

**Affiliations:** The Rockefeller University, New York, NY, USA

## Abstract

What is the cognitive and neural architecture for high-level reasoning? We hypothesize that systems for understanding people and places remain separate throughout the brain, but share a parallel organization. We test this hypothesis using deep neuroimaging of individual human brains on diverse tasks involving reasoning and memory about familiar people, places, and objects. We find that thinking about people and places elicits responses in distinct areas of high-level association cortex, spanning the frontal, parietal, and temporal lobes. Person- and place-preferring brain regions are systematically yoked across cortical zones. These areas have strongly domain-specific response profiles across visual, semantic, and episodic tasks, and are specifically functionally connected to other parts of association cortex with like category preference. Social and spatial networks are anatomically separated even at the top of the cortical hierarchy, and include parts of cortex with anatomical connections to the hippocampal formation. These results demonstrate parallel, domain-specific networks within the cortical apex. They suggest that domain-specific systems for reasoning constitute components of a broader cortico-hippocampal system for long-term memory.

---

How are reasoning systems structured in the human mind and brain? At the cognitive level, some have emphasized the role of domain-general processes that act on different types of information^1,2^. Others have argued for domain-specific systems specialized for learning and reasoning about specific classes of ecologically relevant input, like people, objects, and places^3–5^. Domain-specific neural systems have been most extensively characterized within the realm of perception – in particular, areas of high-level visual cortex that specifically process certain classes of input like faces or scenes^6,7^. In contrast, the domain-specificity of neural systems involved in high-level reasoning and memory remains unclear^8–10^. To what extent are high-level reasoning systems, and higher-order brain regions, specialized for processing certain content domains?

To address this question, we probed functional specialization within zones of human cortex argued to be positioned at the top of a hierarchy of anatomical connectivity^8^. The cortical apex comprises not a single area, but a widely distributed yet interconnected set of brain regions, including medial prefrontal cortex (MPFC), medial parietal cortex (MPC), temporo-parietal junction (TPJ), superior temporal sulcus (STS), and superior frontal gyrus (SFG). Some have argued that this system constitutes a domain-general information processor, integrating diverse types of information from across the brain into abstract representations^8,10,11^. Others have argued for subsystems for distinct cognitive processes, such as social cognition and episodic memory^12,13^, or processing individual concepts and structural relations between concepts^14^.

Here we propose an alternative view: that the cortical apex comprises subsystems for processing information from distinct content domains. We focus on reasoning problems in two ecologically relevant domains: understanding other people, and understanding places or spatial layouts. We hypothesize that parallel subsystems within the cortical apex implement reasoning about people and places separately. In contrast with process-based accounts of functional specialization in association cortex^12,13^, we argue that each of these subsystems supports reasoning, long-term memory, and future prediction, but within distinct content domains.

We evaluate this hypothesis with a human fMRI experiment, using deep imaging of individual human brains, across a range of tasks and domains. To maximize the strength of memory-related responses, we use tasks involving closely familiar people and places for each participant. To test whether responses in association cortex are modulated by content domain consistently across cognitive functions, we used tasks eliciting visual, semantic, and episodic processing. Tasks included visual perception (viewing images of familiar and unfamiliar faces and scenes, and generic objects); semantic judgment (answering questions about personality traits of familiar people, spatial properties of familiar places, and physical properties of generic objects); and episodic simulation (imagining familiar people talking about common conversation topics, navigating through familiar places, and physical interactions of generic objects; Fig 1A, S1). We also performed localizer scans for mental state reasoning or theory of mind (ToM), language comprehension, and dynamic visual perception. Optimized data acquisition and preprocessing methods yielded data with high temporal signal to noise ratio (tSNR, mean 158) despite minimal spatial smoothing, including in regions of signal loss such as the anterior medial temporal lobes (Fig S2).

**Figure 1:**
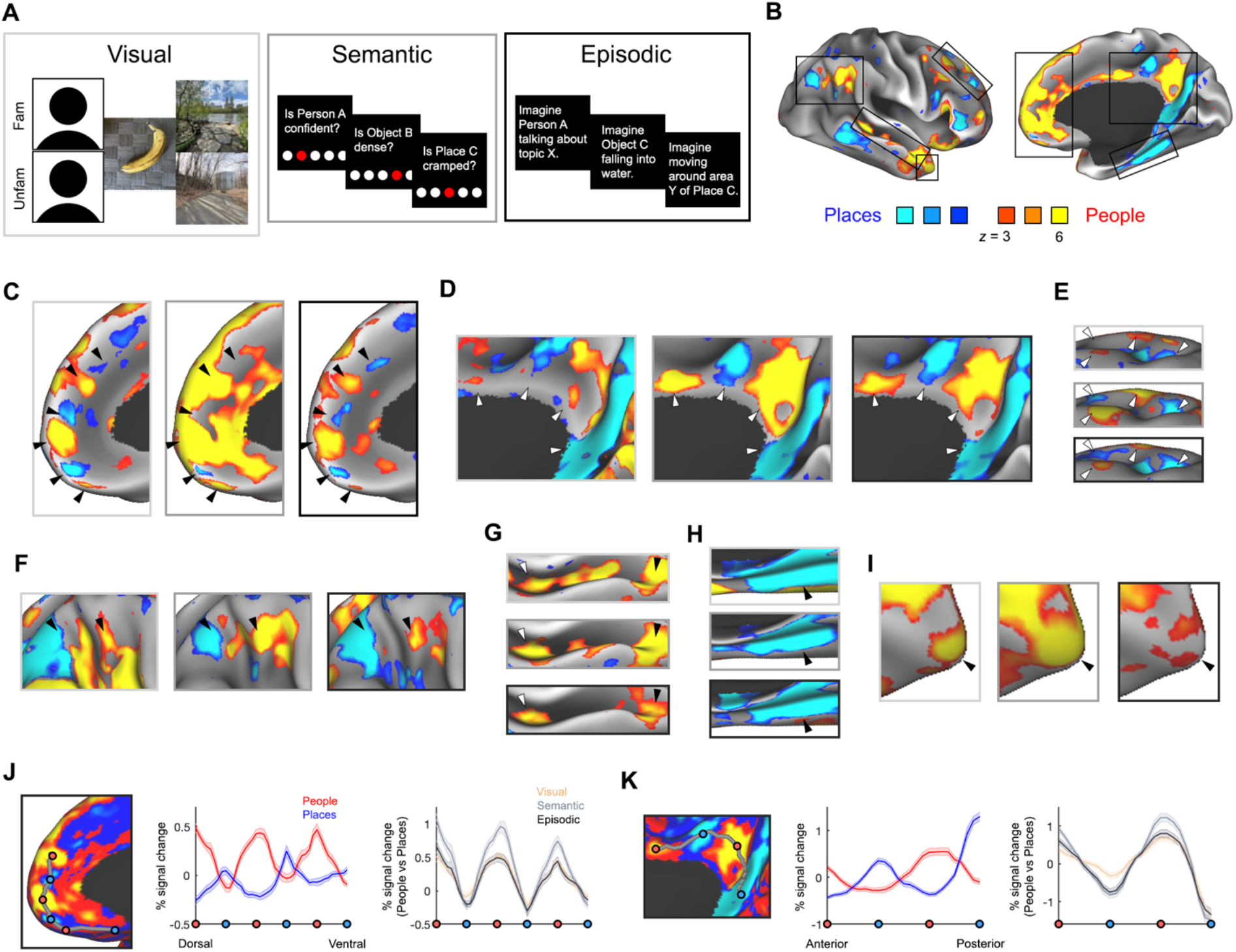
Parallel, distributed responses to people and places across high-level association cortex. **A)** Schematic of conditions from visual, semantic, and episodic tasks. **B)** Whole-brain general linear model-based responses to people versus places, from one representative participant (semantic task, thresholded at a False Discovery Rate of *q* < .01 to correct for multiple comparisons across coordinates). **C)** Responses across each task within medial prefrontal cortex, **D)** medial parietal cortex, **E)** superior frontal gyrus, **F)** temporo-parietal junction, **G)** superior temporal sulcus, **H)** parahippocampal cortex, and **I)** temporal pole. **J)** Analysis of alternating responses to people and places in medial prefrontal cortex. Left: anchor coordinates identified in data half 1. Middle: responses to people and places, averaged across tasks, extracted along cortical geodesics between anchor coordinates in data half 2. Right: responses to people versus places comparison for each task separately. **K)** Analysis of alternating responses to people and places in medial parietal cortex.

We first tested the prediction that social and spatial reasoning engage distinct regions across high-level association cortex. Comparing responses to people and places, we observed preferential responses for both domains among multiple zones distributed across association cortex, including MPFC, MPC, TPJ, and SFG, bilaterally (whole-brain general linear modelbased analysis, corrected for temporal autocorrelation using prewhitening with an ARMA(1,1) model, and corrected for multiple comparisons across coordinates using a false discovery rate of *q* < .01; Figs 1B-F, S6-11). Responses to people were additionally observed within middle and anterior regions of the STS. Importantly, a common pattern of preferential responses to people and places was observed across visual, semantic, and episodic tasks, despite substantial differences in the physical nature of the stimuli (images and words) and cognitive demands of the tasks. These preferences thus cannot be explained by a confound specific to any one task, and are more parsimoniously explained as effects of content domain.

We next tested the prediction that specializations for places and people are not simply segregated, but organized in parallel. Across multiple zones of association cortex, we found that regions responsive to people and places were systematically yoked, with adjacent or alternating parts of cortex showing opposing stimulus preferences. In the TPJ, a pair of regions was typically observed, with a person-preferring area just anterior to a place-preferring area (Fig 1F). In MPFC and MPC, a series of interdigitated responses to people and places was observed along a curved axis following the cingulate sulcus, with 3-5 (MPFC) or 2-3 (MPC) pairs of regions in individual participants (Fig 1C-D). To formally analyze this pattern of alternation, we performed a split-half analysis. In data from even-numbered runs, we used whole-brain statistical maps, combined across tasks, to identify peak coordinates (anchors) of the alternating response to people and places in each participant and hemisphere. We then extracted responses (% signal change) to people and places in left-out data from odd runs, along a path defined by geodesics on the cortical surface between anchors. This analysis verified an alternating pattern of response to people and places in medial prefrontal and parietal cortex, consistent across tasks (Fig 1J-K). These results demonstrate that systems for processing people and places share a parallel anatomical organization across association cortex.

The cortical apex is positioned far, in terms of connection distance, from the sensory periphery and near the hippocampus and limbic system^8^. We thus wondered whether person and place preferences exist in zones of cortex known to have direct anatomical connections with the hippocampal formation. We first tested for the presence of place responses in parahippocampal cortex (PHC) and retrosplenial cortex (RSC), areas with bidirectional connections to the hippocampus via entorhinal cortex and the subicular complex^15–17^, implicated in spatial cognition in rodents and primates^18,19^. Whole-brain analyses showed place responses in PHC and RSC, defined using the Human Connectome Project’s multimodal cortical surface atlas^20^, which were consistent across tasks and participants (left hemisphere, 60/60 comparisons; right hemisphere, 60/60 comparisons; Figs 1H, S6-8). We then tested for the presence of person responses in the temporal pole (TP), a primate-specific brain region with bidirectional connections to the hippocampus via entorhinal cortex^15,21^. A region of TP specialized for social cognition has long been hypothesized, but has not been identified reliably or in individual brains^22^. We observed a region within TP that responds preferentially to people over places, consistent across tasks and participants (left hemisphere, 30/30 comparisons; right hemisphere, 29/30 comparisons; Figs 1I, S6-8). These results demonstrate the presence of person and place preferences within parts of cortex with anatomical connections to the hippocampal formation in primates, identifying putative relay areas between category-preferring regions of association cortex and the hippocampus.

Our results thus far have demonstrated that regions with preferential responses to people and places have a parallel, distributed organization across association cortex. To what extent are these regions specialized for processing information from their preferred content domain? To address this question, we assessed the magnitude of responses across content domains and tasks. Functional regions-of-interest (ROIs) were defined as the top 5% of maximally person- or placepreferring coordinates (semantic task) within anatomical search spaces covering zones of the cortical apex: MPC, MPFC, TPJ, STS, SFG, TP, and PHC (Fig 2A-B). We then extracted response magnitudes from independent data, across all task conditions.

**Figure 2:**
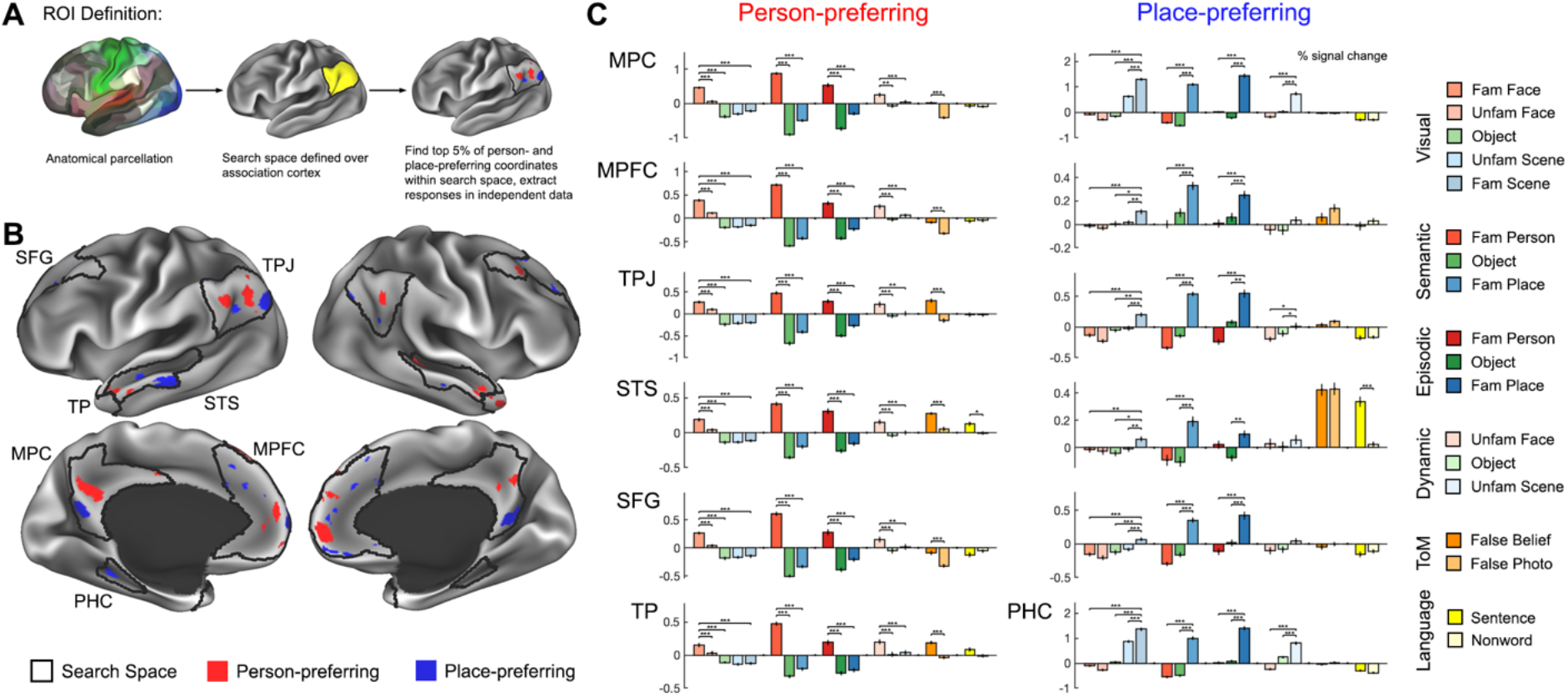
Region-of-interest (ROI)-based analysis reveals domain-specific responses across tasks. **A)** ROIs were defined as the top 5% of person- or place-preferring coordinates within anatomical search spaces. **B)** Search spaces and example functional ROIs from one representative participant. **C)** Responses (% signal change) extracted from functionally defined ROIs, across all task conditions. Error bars show standard error across runs. * *P* < .05/7 = .0071, ** *P* < 10^-3^, *** *P* < 10 ^-4^ (linear mixed model across runs, with participant included as random effect). Abbreviations: MPC, medial parietal cortex; MPFC, medial prefrontal cortex; TPJ, temporo-parietal junction; STS, superior temporal sulcus; SFG, superior frontal gyrus; TP, temporal pole; PHC, parahippocampal cortex.

This ROI analysis found that responses in category-sensitive subregions of association cortex were strongly selective for their preferred stimulus category (Fig 2C). Across ROIs, responses to non-preferred stimulus categories were typically at or below baseline. Personpreferring areas in MPC, MPFC, TPJ, STS, SFG, and TP responded significantly more strongly to seeing images of familiar people versus objects and familiar scenes; making judgments about familiar people versus objects and familiar places; and imagining events involving familiar people versus objects and familiar places (all 36 *P*’s < .0071 = .05 / 7, applying Bonferroni correction across the full set of 7 ROIs; individual values in Table S3; linear mixed effects model across runs, with participant included as a random effect). Similarly, place-preferring areas in MPC, MPFC, TPJ, STS, SFG, and PHC responded significantly more strongly to seeing images of familiar scenes versus objects and familiar people; making judgments about familiar places versus objects and familiar people; and imagining events involved familiar places versus objects and familiar people (all 36 *P*’s < .0071; individual values in Table S4). Strong category selectivity was not only observed for the focal, maximally responsive areas used in this analysis, but across a range of ROI sizes, from the top 5-40% of category-preferring coordinates (Fig S12). Other regions of association cortex responded selectively to object or language conditions (Figs S13-14, Tables S5-6), arguing against an explanation of person or place responses in terms of generic factors like attention or task engagement. These results demonstrate domain-specificity as a strong organizing principle of association cortex, arguing for distinct, specialized systems for social and spatial reasoning.

Given its anatomical positioning near the medial temporal lobe, the cortical apex is well-positioned to play a role in long-term memory. How do person- and place-preferring regions respond when processing familiar and unfamiliar entities? Person-preferring areas in MPC, MPFC, TPJ, STS, SFG, and TP responded significantly more strongly to familiar versus unfamiliar face images (all 6 *P*’s < .0071; Table S3). Similarly, place-preferring areas in MPC, MPFC, TPJ, STS, SFG, and PHC responded significantly more strongly to familiar versus unfamiliar scene images (all 6 *P*’s < .0071). These complementary familiarity effects suggest distinct neural systems supporting long-term memory for social and spatial information.

Does the brain use common or distinct systems for abstract reasoning about others’ mental states (ToM), and storing information in long-term memory about specific familiar people (social memory, also termed person knowledge)? While ToM and social memory are often described as separate processes, another possibility – consistent with our argument for functional specialization by content domain – is that these concepts reflect different aspects of a common cognitive process. For example, social memory could consist of ToM-like internal models for familiar people. In contrast, theories arguing for functional specialization based on a process distinction between social reasoning and long-term memory predict separate neural substrates for ToM and social memory.

To address this question, we asked whether brain regions responsive to familiar person tasks also respond when participants reason about the mental states of unfamiliar characters in a story. Person-preferring areas in MPC, MPFC, TPJ, STS, SFG, and TP responded significantly more strongly during reasoning about false beliefs relative to false “photographs” or physical representations (all 6 *P*’s < .0004 = .05 / 12, applying Bonferroni correction across the full set of person- and place-preferring ROIs; Fig 2C, Table S3). Conversely, ROIs in MPC, MPFC, TPJ, STS, SFG, and TP defined by the ToM localizer responded significantly more strongly to familiar people over objects and familiar places, across visual, semantic, and episodic tasks (all 36 *P*’s < .0071; Table S7). In contrast, place-preferring areas in MPC, MPFC, TPJ, STS, SFG, and PHC did not respond more strongly to false beliefs over photographs (all 6 *P*’s > .9; Table S4). These results argue for a common system for ToM and social memory. They support our hypothesis of a set of regions with domain-specific responses to social content across a range of tasks.

How are functionally specialized regions of association cortex situated along a connectivity-based cortical hierarchy? We hypothesize that person- and place-preferring brain regions extend to the top of the cortical hierarchy. To operationally define the position of brain regions in a connectivity-based hierarchy, we computed low-dimensional embeddings of graphs defined by resting-state correlation distance in individual participants, using the diffusion maps algorithm to effectively capture both local and global distance structure^8,23^. Consistent with prior results from group-level data, the space spanned by the first two principal dimensions included an “apex gradient” separating unimodal sensorimotor cortices from a transmodal cortical apex, along with a “visuomotor gradient” separating visual from somatomotor and auditory cortices (Fig 3A). For visualization purposes, a boundary was placed at an arbitrary position along the apex gradient, separating areas roughly considered within versus outside of the cortical apex.

**Figure 3:**
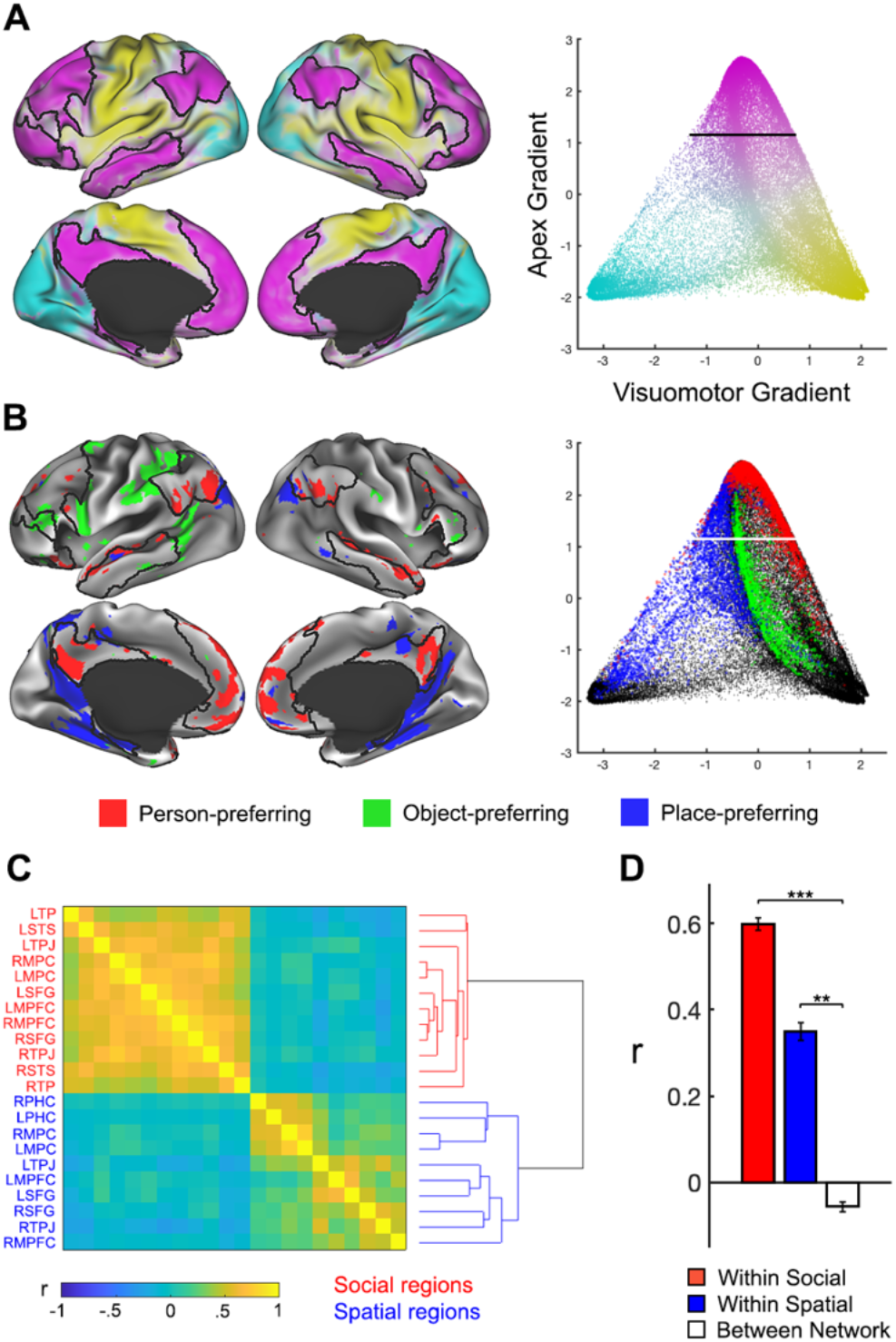
Person- and place-preferring areas form functionally coupled networks within the cortical apex. **A)** Diffusion embedding of resting-state functional connectivity data reveals large-scale cortical gradients, separate the cortical apex (pink) from visual (cyan) and somatomotor (yellow) cortices. **B)** Person-, place-, and object-preferring regions (coordinate-wise *Z* > 2.3 across visual, semantic, and episodic tasks), shown on the cortical surface and in the diffusion embedding. All results from one representative participant. **C)** Matrix of resting-state correlations among person- and place-preferring areas, and hierarchical clustering results. **D)** Mean correlations within social and spatial networks, and between the two. Error bars show standard error across correlation values. ** *P* < 10^-3^, *** *P* < 10^-4^ (permutation test). Abbreviations: MPC, medial parietal cortex; MPFC, medial prefrontal cortex; TPJ, temporo-parietal junction; STS, superior temporal sulcus; SFG, superior frontal gyrus; TP, temporal pole; PHC, parahippocampal cortex.

We next identified person-, place-, and object-preferring coordinates in this space, only including coordinates that responded significantly across visual, semantic, and episodic tasks (Fig 3B, S16; *Z* > 2.3 for each task). The three sets of regions, while distributed and interdigitated on the cortical surface (as shown before, Fig 1), were each clustered in connectivity distance space. Person-preferring regions were positioned at or near the cortical apex. Place-preferring areas spanned nearly the whole length of the apex gradient, stretching from visual cortex up to the cortical apex. This is consistent with prior findings associating place areas with multiple functional networks^24,25^. On the cortical surface, place responses often straddled the boundary between visual and apex cortex, in medial and lateral parietal cortex as well as ventral temporal cortex. By contrast, object-preferring areas were typically clustered toward the middle or low end of the principal gradient, adjacent to somatomotor cortex. These results demonstrate that person- and place-preferring regions of cortex both exist within the cortical apex. The results also show that they are not equal: place-preferring regions extend “lower” into visual cortex than person-preferring ones.

To what extent do person- and place-preferring regions across multiple zones of cortex constitute functionally coupled networks, rather than sets of distinct, isolated processors? While the results shown in Figure 3B indicate that areas with common stimulus preferences also share patterns of functional connectivity, we next tested this hypothesis explicitly using the ROIs defined above (Fig 2B). We found that resting-state correlations were substantially stronger within person- or place-preferring areas than between the two (Fig 3C-D; permutation test; *P* < 10^-4^, person vs between; *P* < 10^-3^, place vs between). Hierarchical clustering of regions based on correlation distance revealed a dominant two-cluster solution, separating person- and placepreferring regions (Fig 3C). Whole-brain functional connectivity analyses using person- and place-preferring regions as seeds confirmed this pattern of results, showing resting-state correlations to other parts of the frontal, temporal, and parietal lobes with similar stimulus preferences (Figs S17-20). These results justify the description of these sets of areas as social and spatial “networks” – functionally coupled sets of brain areas distributed across cortex, with common stimulus selectivity.

Taken as a whole, these results demonstrate that social and spatial reasoning elicit domain-specific responses in distributed networks with parallel anatomical organization (Fig 4A). Similarity between patterns of response profile and functional connectivity among person- and place-preferring regions from different parts of high-level association cortex – the medial and lateral frontal, parietal, and temporal lobes – argues for two coherent networks. Yoked responses to people and places within common cortical zones indicate separate but parallel systems. These findings suggest against the characterization of the cortical apex as a domain-general integrator of information from across the brain^8^, instead demonstrating that domain-specific responses extend beyond visual cortex all the way to the top of the cortical hierarchy. Consistent responses to people and places across visual, semantic, and episodic tasks provide strong evidence for a functional dissociation driven by content domain, rather than cognitive process^12,13^.

**Figure 4:**
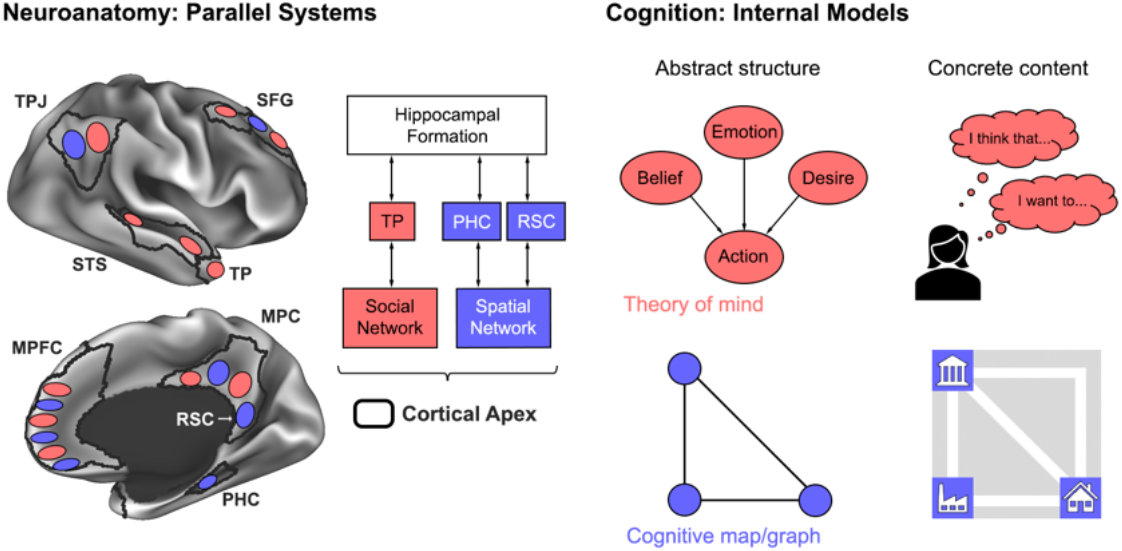
Theoretical framework – parallel systems for social and spatial reasoning. Left: Social and spatial reasoning are implemented by parallel, distributed networks of association cortex. Each network is positioned within the cortical apex and interacts closely with the hippocampal formation. Right: We propose that these systems support a common cognitive operation within distinct domains, forming internal models of familiar people and places by combining abstract structural relations (theories of mind and cognitive maps or graphs) with concrete information from experience.

Our results identify a novel principle of cortical functional organization – yoked responses to people and places – involving a combination of newly discovered and previously identified brain regions. We characterize a socially selective region of the temporal pole, hypothesized but not reliably detected in prior studies^22^. We identify novel place-preferring regions of SFG and MPFC, consistent with a role for medial frontal cortex in place understanding that has been hypothesized^26^ and supported by lesion evidence^27^. Our work also provides new insight into the function of previously identified brain regions involved in social reasoning^9,12,28^ and place understanding^25,29^. By studying these regions using deep imaging of individual humans across a wide range of tasks, we provide a precise characterization of their functional organization, identify relationships between social and spatial domains, and provide strong evidence for domain-specificity across multiple tasks and stimulus classes, beyond what has been possible so far.

Why might the brain employ parallel mechanisms for understanding people and places? While these two problems may appear different on their face, they share a similar structure. In navigating the physical world, we decide how to move around in an uncertain, constantly changing environment, in order to execute plans and optimize rewards. In navigating the social world, we decide what to say and do around people with uncertain, constantly changing internal states, in order to achieve personal and collective goals. In both cases, researchers have argued that the mind solves these problems using abstract, internal models – theories of mind^30–32^, and cognitive maps or graphs^33,34^ – combined with concrete information about specific familiar people and places derived from experience^31,35,36^.

We propose that social and spatial networks support the common cognitive operation of building internal relational models, but within distinct content domains (Fig 4B). These models can in turn be used for structuring long-term memories, reasoning about events in the present, and predicting what will happen in the future. In contrast with classic cognitive psychological theories – which posit a division of labor between processes such as reasoning, decision-making, and long-term memory – we argue that a common mechanism supports these apparently diverse processes. This view provides a parsimonious explanation of the domain-specific responses we observed across a range of tasks, including visual perception, semantic judgment, episodic recall, and ToM reasoning.

Social and spatial networks extended to the apex of a connectivity-based cortical hierarchy, and included input zones to the hippocampal formation, supporting the hypothesis that these networks are anatomically positioned near the hippocampus (Fig 4). This indicates that the neural system supporting human social reasoning can be understood as a domain-specific component of a broader corticohippocampal long-term memory system^37,38^. In line with our theoretical framework, this system has been argued to construct flexible models of the external world by establishing conjunctions between abstract structural relations and concrete content^14,39^. While this system has been studied extensively in the domain of spatial navigation, our work indicates that a similar mechanism is used for understanding people, and identifies a region of TP that is anatomically well-positioned to support interactions between the social network and hippocampal formation.

Our empirical results establish a new approach for addressing the mystery of how humans understand other people. Just as interactions between the hippocampus and functionally specialized cortical areas have been argued to underly the ability to learn cognitive maps of the spatial environment^19,34^, we propose that interactions between the hippocampus and a socially specialized network support learning causal models of specific familiar people. This work points to numerous directions for future research, into how the brain builds representations of familiar people during learning, and how networks for social and spatial reasoning emerged in evolution and differentiate in development.

## Methods

### Participants

Ten human participants (5 male, 5 female; age 28-40) were scanned using fMRI. Participants were healthy with normal or corrected vision, right-handed, and native English speakers. The experimental protocol was approved by the Rockefeller University Institutional Review Board, and participants provided written, informed consent.

### Tasks

We used a range of perceptual and cognitive tasks involving familiar people and places. Participants were asked to choose six of their top ten most familiar people and places, and tasks involved processing these six people and places. Main tasks included: visual perception of familiar and unfamiliar faces and scenes, and generic objects; semantic judgment about familiar people and places, and generic objects; and episodic simulation of events involving familiar people and places, and generic objects (Fig 1A, S1). Additionally, localizer tasks from the existing literature were run, including tasks eliciting theory of mind (ToM; reading and answering questions about stories involving false beliefs or false physical representations)^40^; language comprehension (reading sentences or nonword lists)^41^; and dynamic visual perception (watching videos of moving faces, objects, and scenes)^42^. In each of three scans, one main task and one localizer were run (order: visual and ToM, semantic and dynamic perception, episodic and language). Stimulus presentation scripts can be found at https://osf.io/5yjgh/.

In the visual perception task, participants viewed serially presented naturalistic images of faces, objects, and scenes. For each participant, we obtained 20 images each of six familiar faces and scenes. Familiar face images were obtained directly from friends or family members, without the participant seeing them. Control images were defined as six yoked unfamiliar faces and scenes. Familiar and unfamiliar faces were matched on age group (young adult, middle-aged, old), race, and gender; familiar and unfamiliar scenes were matched on rough semantic category (e.g. outdoor street view; building interior). Object images were of six generic objects with varying physical properties - a banana, a baseball, a feather, a rock, a sponge, and a wrench. Face and object images contained no clear spatial structure (e.g. corners), and had minimal contextual cues beyond the background. Scene images contained no people. All five image categories were matched, for each participant, on the mean and variance across images of luminance, root-meansquare contrast, and saturation (in CIE-Lch space, with a D65 illuminant): all *P*’s > .05, one-way ANOVA and Bartlett test. Images were presented at 768 x 768 resolution, 12.8 x 12.8 degrees of visual angle, for 1.85s each with a 150ms interstimulus interval, in 18s blocks of images of one identity. Participants performed a one-back task, pressing a button when an image was repeated. A post-scan questionnaire verified that participants could recognize the person or place in the familiar images (mean: 100% for faces, 84% for scenes), but not the unfamiliar images (mean: 0% for faces, 2% for scenes, Fig S4).

In the semantic task, participants rated traits of familiar people and places, and generic objects, on a 0 to 4 scale. This included personality traits of people (e.g. confident, angry, intelligent), spatial or navigational properties of places (e.g. cramped, large, has walls), and physical properties of objects (e.g. soft, heavy, rough; Table S1). Participants rated by moving an icon left or right, over 18s blocks of four questions for a given identity, for a total of 20 questions per condition and identity.

Prior to the episodic task, participants listed five common conversation topics for each familiar person, and five familiar subregions of each familiar place. In the scan, they were asked to imagine familiar people talking about common topics, navigating specific subregions of familiar places, and objects engaged in physical interactions (e.g. rolling or sliding down a hill, falling into water). Participants were specifically asked to generate a novel event, rather than remember a past event. Imagination blocks lasted 18s, with a 3s verbal prompt, and a 1s hand icon at the end of the block, which participants responded to with a button press. After the scan, participants were asked (for 2/5 of trials) questions about difficulty, detail, visual imagery, emotional response, and memory recall (Fig S5). Additionally, a free response description was collected, to ensure that participants could describe what they imagined. 9/10 participants were able to describe all events; the remaining one recalled 75% of events.

Across the three main tasks, blocks were separated by 4s of resting fixation, and presented in five 8-13 minute runs per task, with palindromic block orders, counterbalanced across runs and participants. Fixation blocks were included in the beginning, middle, and end of the experiment to estimate a resting baseline. Localizer tasks were split into 4-5 minute runs, with four runs for theory of mind and language tasks, and six for dynamic perception. Scanner task performance was high, indicating sustained attention during scan sessions (Fig S3). After each scan, participants were asked to rate on a 0-10 scale to what extent they recalled memories of events during the person and place conditions (Fig S4).

### Behavioral data analysis

To evaluate participants’ self-reported experience in the episodic task, we compared behavioral responses across each pair of conditions (person, object, place), for each of the five questions. Statistics were computed using a linear mixed effects model (MATLAB’s fitlme, unpaired, two-tailed comparisons) across items or event ratings, with random intercepts for participant.

### MRI acquisition

Participants were scanned on a Siemens 3T Prisma across three 2.5-hour sessions, which included task acquisitions as well as 40 minutes of high-resolution anatomical images, 60 minutes of resting-state acquisitions, and spin echo acquisitions for distortion correction. Three each of T1- and T2-weighted anatomical images were acquired at 8mm resolution. Task and resting-state data were acquired using a multiband, multi-echo EPI pulse sequence, optimized to boost temporal signal-to-noise ratio (tSNR) throughout the brain and specifically in the anterior medial temporal lobes (TR = 2s, TE = 14.4, 33.9, 53.4, 72.9, and 92.4ms, 2.4×2.4×2.5mm resolution, 48 oblique axial slices with near whole brain coverage, multiband acceleration 3x, GRAPPA acceleration 2x, interleaved slice acquisition). 3-4 parameter-matched spin echo acquisitions were acquired per scan, between every four runs of task. Raw MRI data can be found at https://openneuro.org/datasets/ds003814.

### MRI preprocessing

Data were preprocessed and analyzed using a custom pipeline, integrating software elements from multiple software packages: FSL (6.0.3), Freesurfer (7.1.1), AFNI, Connectome Workbench 1.5, tedana 0.0.10, and Multimodal Surface Matching (MSM). The code is available at https://github.com/bmdeen/fmriPermPipe/releases/tag/v2.0.1, with datasetspecific wrapper scripts at https://github.com/bmdeen/identAnalysis.

Anatomical images were preprocessed using an approach based on the Human Connectome Project (HCP) pipeline^43^. The three images for each modality were linearly registered using FLIRT^44^ and averaged; registered from T2- to T1-weighted; aligned to ACPC orientation using a rigid-body registration to MNI152NLin6Asym space; and bias-corrected using the sqrt(T1*T2) image^45^. Cortical surface reconstructions and subcortical parcellations were generated using Freesurfer’s recon-all^46^. Surface-based registration (MSMSulc) was used to register individual surfaces to fsLR average space^47^. This registration was used to project the HCP multimodal cortical parcellation^20^ onto individual surfaces.

Functional data were preprocessed using a pipeline tailored to multi-echo data, aiming to optimize tSNR while maintaining high spatial resolution. Motion parameters were first estimated using MCFLIRT^48^. Intensity outliers were removed using AFNI’s 3dDespike, and slice timing correction was performed using FSL’s slicetimer. Motion correction was then applied, in combination with topup-based distortion correction^49^, and rigid registration to a functional template image, in a single-shot transformation with one linear interpolation, to minimize spatial blurring. Multi-echo ICA was performed using tedana, with manual adjustments, to remove nonblood-oxygen-dependent (non-BOLD) noise components^50^. Data were intensity normalized and resampled to an individual-specific CIFTI space aligned with the anatomical template image, with 32k density fsLR surface coordinates, and 2mm volumetric subcortical coordinates. Registration between the functional and anatomical templates was computed using boundarybased registration (bbregister)^51^. Surface data and subcortical volumetric data were both smoothed with a 2mm-FWHM Gaussian kernel. For resting-state data, the global mean signal was removed via linear regression, to diminish global respiratory artifacts not removed by multiecho ICA^52^.

### Whole-brain analysis

Whole-brain statistical analyses were performed in individuals using AFNI’s 3dREMLfit, modeling temporal autocorrelation with a coordinate-wise ARMA(1,1) model^53^. Results were thresholded using a false discovery rate of *q* < .01 (two-tailed), to correct for multiple comparisons across ~90K CIFTI coordinates^54^. We compared responses to people versus places, and objects versus places, across tasks (boxcar regressors convolved with canonical double gamma hemodynamic response function). To determine anatomical locations of responses, we used the HCP multimodal cortical parcellation, with PHC defined as areas PHA1-3, RSC defined as area RSC, and TP defined as area TGd.

### Interdigitation analysis

To test for spatially interdigitated responses, we used part of the dataset (even runs), combined across visual, semantic, and episodic tasks, to identify person- and place-sensitive parts of medial prefrontal and parietal cortex. We identified 3 pairs of regions in medial prefrontal cortex, and 2 pairs in medial parietal cortex, in each participant and hemisphere, and hand-drew anchor coordinates over these responses. We then defined paths corresponding to geodesics on the cortical surface between anchors, and extracted person and place responses (% signal change) in left-out data from odd runs. To combine measurements across participants, we interpolated responses along each path to a standardized coordinate frame with 7 equidistant coordinates between each anchor.

### Region-of-interest (ROI) analysis

ROI-based analyses were conducted to assess how person- and place-preferring brain areas respond across a range of task conditions. ROIs were defined as the top 5% of person- or place-preferring coordinates (semantic task) within anatomical search spaces capturing zones within the cortical apex: medial frontal cortex, medial parietal cortex, temporo-parietal junction, superior frontal gyrus, superior temporal sulcus, parahippocampal cortex, and temporal pole. Search spaces were composed of regions from the HCP parcellation (Table S2). To extract responses to the task used to define ROIs, a leave-one-run-out analysis was performed, in which ROIs were defined in all but one run of data, and responses extracted in the left out run. Statistics were performed on percent signal change values across runs, using a linear mixed effects model (MATLAB’s fitlme), with participant included as a random effect. Similar analyses were performed for object-sensitive regions, and for the language and theory of mind localizers. Analyses were also performed for a range of ROI sizes, from 5 to 40% (Fig S12).

For ROIs defined as person-preferring or by the language or ToM tasks, responses to people were compared with places and objects, across visual, semantic, episodic, and dynamic tasks. Additionally, responses to familiar/unfamiliar faces were compared. For place-preferring ROIs, responses to places were compared with people and objects across tasks, and responses to familiar/unfamiliar faces were compared. For object-preferring ROIs, responses to objects were compared with people and places, across tasks. For all ROIs, we compared responses to false belief versus false photo and sentence versus nonword list conditions. All comparisons were paired and one-tailed.

### Connectivity gradient analysis

To position functional responses along large-scale connectional gradients in cortex, we computed a low-dimensional embedding of resting-state functional connectivity similarity, using the diffusion maps algorithm^23^. Following Margulies et al.^8^, we computed a pairwise correlation matrix between resting-state time series from ~59K cortical surface coordinates, kept only the top 10% of values per row, zeroed negative values, and computed cosine similarity between rows to generate a positive, symmetric similarity matrix. We then generated a diffusion map embedding, using α = .5 (Fokker-Planck normalization), and automated estimation of diffusion time via spectral regularization, by multiplying Laplacian eigenvalues *λ_i_* by 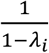. Across participants, the space spanned by the first two principal directions included previously described anatomical gradients^8^: one between sensory cortex and transmodal association cortex (apex gradient), and one between sensorimotor and visual cortex (visuomotor gradient, Fig 4). In order to align the apex gradient with the y-axis, a manual rotation of 15° was applied to this space for visualization. Because the orientation of the apex and visuomotor gradients in diffusion embedding space varied across participants, a Procrustes transformation was used to map each participant’s embedding to the first participant’s, using the top three dimensions. To overlay functional responses on the diffusion embedding, we identified coordinates that were responsive to either 1) people versus objects and people, 2) places versus people and objects, or 3) objects versus people and places, across visual, semantic, and episodic tasks (coordinate-wise *Z* > 2.3 for each task). These coordinates were displayed in color in diffusion space. Only a small proportion (< 0.15% for each participant) of coordinates were responsive to multiple contrasts. These were colored with the following precedence: object-preferring, place-preferring, person-preferring. For the sake of visually comparing functional responses to the apex gradient, boundaries were drawn on the cortical surface at the 70^th^ percentile of coordinates along the apex gradient, using a smoothed map of gradient values (surface-based 8mm-FWHM Gaussian).

### ROI-based functional connectivity analysis

Resting-state correlations were computed between person- and place-preferring areas defined similarly to the ROI analysis, but using separate search spaces for each hemisphere. Place-preferring STS subregions were not included, because such responses were not reliably observed across participants. The regions were hierarchically clustered based on correlation distance, and regions were ordered in a way that minimized distance between adjacent pairs, without separating clusters (Fig 4A; MATLAB’s optimalleaforder). Within-versus between-network correlations were compared using a permutation test, permuting regions (10,000 iterations). Whole-brain correlation maps, with functional ROIs as seeds, were computed for person- and place-preferring regions of left MPC and SFG, within each individual participant. These two regions were chosen for presentation because the functional connectivity of other ROIs generally resembled one of these two.

## Data and code availability

Raw data are available at https://openneuro.org/datasets/ds003814. Stimulus materials are available at https://osf.io/5yjgh/. Analysis code is available at https://github.com/bmdeen/fmriPermPipe/releases/tag/v2.0.2 (generic analysis tools) and https://github.com/bmdeen/identAnalysis (dataset-specific wrapper scripts).

## Acknowledgements

We thank the staff at the Cornell Citigroup Biomedical Imaging Center for assistance with data acquisition, and Charles Lynch for helpful discussion on pulse sequence design. This work was supported by fellowships from the Helen Hay Whitney and Leon Levy foundations (B.D.), and the Center for Brains, Minds & Machines, funded by National Science Foundation STC award CCF-1231216 (W.A.F.).

## Author Contributions

B.D. and W.A.F. conceived and designed the experiment. B.D. collected and analyzed the data. B.D. and W.A.F. wrote the paper.

