## Supplementary Info for "Parallel systems for social and spatial reasoning within the cortical apex"

| **People** |  | **Objects** |  | **Places** |  |
| --- | --- | --- | --- | --- | --- |
| nice | mean | soft | hard | large | small |
| trustworthy | greedy | rough | smooth | spacious | cramped |
| caring | cold | light | heavy | narrow | wide |
| rational | emotional | warm | cold | underground | above ground |
| incompetent | smart | fragile | unbreakable | exposed | skyscraping |
| determined | fearful | round | angular | flat | gated |
| lazy | serious | rigid | bendable | has many entrances | has rooms |
| extroverted | shy | dense | bouncy | has multiple levels | has specific pathways |
| aggressive | angry | sticky | fuzzy | has a roof | has high ceilings |
| anxious | calm | bumpy | squishy | has walls | has ramps |

**Table S1.** Full set of adjectives using for semantic task, with 20 adjectives per condition.

| **Search Space** | **Parcels** |
| --- | --- |
| MPC | RSC, POS1, v23ab, 7m, PCV, 31pv, 31pd, d23ab, 31a, 23d |
| MPFC | 8BM, a32pr, p24, d32, 9m, a24, p32, 10d, 25, s32, 10r, 10v |
| TPJ | PFm, PGi, PGs |
| STS | STSda, STSva, TE1a, STSdp, STSvp, TE1m |
| SFG | 9a, 9p, 8BL, 8Ad, 8Av |
| TP | TGd |
| PHC | PHA1, PHA2, PHA3 |
| Premotor | 6r |
| IFS | IFSa |
| PF | PF, PFop, PFt |
| PHT | PHT |
| AntTemp | STSda, STSva, TE1a, TE2a, STGa, TGd |
| PostTemp | TPOJ1, STV, STSdp, STSvp, PHT, TE1p, TE1m |
| IFG | IFJa, IFSp, 44, 45, 47l |
| MFG | FEF, 55b |

**Table S2.** How search spaces were defined in terms of anatomical parcels from the multimodal parcellation (Glasser et al. 2016).

|  | **MPC** | **MPFC** | **TPJ** | **STS** | **SFG** | **TP** |
| --- | --- | --- | --- | --- | --- | --- |
| Person vs Place (Visual) | 3.30E-12 | 2.20E-16 | 1.10E-09 | 2.10E-14 | 8.80E-11 | 2.60E-06 |
| Person vs Object (Visual) | 3.20E-23 | 8.50E-23 | 3.40E-11 | 4.20E-18 | 6.70E-16 | 4.10E-06 |
| Familiar vs Unfamiliar Face | 2.20E-16 | 3.80E-13 | 5.40E-07 | 1.60E-08 | 7.00E-07 | 2.70E-06 |
| Person vs Place (Semantic) | 1.60E-15 | 8.50E-23 | 7.80E-12 | 8.90E-16 | 1.10E-11 | 2.00E-06 |
| Person vs Object (Semantic) | 3.20E-23 | 8.50E-23 | 2.20E-13 | 4.20E-18 | 1.10E-13 | 2.00E-06 |
| Person vs Place (Episodic) | 2.10E-10 | 3.20E-09 | 2.50E-10 | 9.70E-09 | 3.60E-08 | 4.20E-08 |
| Person vs Object (Episodic) | 6.50E-13 | 5.00E-12 | 5.60E-12 | 6.10E-10 | 2.40E-08 | 5.50E-08 |
| Person vs Place (Dynamic) | 4.50E-05 | 1.60E-07 | 2.90E-04 | 1.70E-06 | 3.70E-04 | 3.90E-08 |
| Person vs Object (Dynamic) | 4.50E-04 | 2.50E-10 | 5.50E-07 | 7.20E-08 | 1.30E-06 | 9.50E-10 |
| Belief vs Photo | 1.00E-07 | 3.50E-06 | 6.20E-08 | 5.30E-09 | 1.10E-07 | 4.70E-08 |
| Sentences vs Nonwords | 8.10E-01 | 1.00 | 9.40E-01 | 4.50E-03 | 1.00 | 1.10E-02 |

**Table S3.** *P*-values from person-preferring ROI analysis. Linear mixed model across runs, with participant included as random effect.

|  | **MPC** | **MPFC** | **TPJ** | **STS** | **SFG** | **TP** |
| --- | --- | --- | --- | --- | --- | --- |
| Place vs Person (Visual) | 2.00E-15 | 5.00E-07 | 9.10E-09 | 4.50E-04 | 9.40E-08 | 4.40E-16 |
| Place vs Object (Visual) | 1.90E-02 | 3.00E-03 | 7.90E-04 | 3.30E-03 | 4.60E-06 | 2.40E-13 |
| Familiar vs Unfamiliar Scene | 8.00E-15 | 4.70E-04 | 1.40E-07 | 1.30E-04 | 1.80E-05 | 1.10E-09 |
| Place vs Person (Semantic) | 1.90E-02 | 1.70E-09 | 2.70E-14 | 4.20E-06 | 1.50E-13 | 9.10E-15 |
| Place vs Object (Semantic) | 1.90E-02 | 2.30E-08 | 1.80E-09 | 9.50E-07 | 1.40E-09 | 6.10E-13 |
| Place vs Person (Episodic) | 2.90E-12 | 1.60E-06 | 2.50E-12 | 1.60E-01 | 3.20E-08 | 2.20E-11 |
| Place vs Object (Episodic) | 1.00E-11 | 4.80E-06 | 1.00E-04 | 1.40E-04 | 1.30E-06 | 8.70E-10 |
| Place vs Person (Dynamic) | 1.00E-08 | 9.80E-03 | 6.80E-03 | 5.10E-01 | 2.80E-02 | 9.90E-09 |
| Place vs Object (Dynamic) | 5.90E-08 | 7.10E-02 | 5.10E-03 | 9.90E-02 | 3.00E-02 | 5.40E-05 |
| Belief vs Photo | 9.00E-01 | 1.00 | 1.00 | 1.00 | 1.00 | 1.00 |
| Sentences vs Nonwords | 1.00 | 1.00 | 1.00 | 2.40E-05 | 1.00 | 2.00E-02 |

**Table S4.** *P*-values from place-preferring ROI analysis. Linear mixed model across runs, with participant included as random effect.

|  | **Premotor** | **IFS** | **PF** | **PHT** |
| --- | --- | --- | --- | --- |
| Object vs Person (Visual) | 7.60E-01 | 6.50E-03 | 1.20E-04 | 9.20E-02 |
| Object vs Place (Visual) | 1.80E-06 | 2.10E-09 | 6.40E-14 | 6.60E-07 |
| Object vs Person (Semantic) | 1.10E-09 | 6.70E-16 | 9.20E-11 | 4.00E-03 |
| Object vs Place (Semantic) | 4.00E-10 | 3.20E-03 | 6.00E-05 | 1.80E-03 |
| Object vs Person (Episodic) | 6.20E-10 | 1.80E-09 | 4.80E-09 | 2.80E-05 |
| Object vs Place (Episodic) | 2.00E-08 | 5.10E-10 | 1.00E-09 | 3.30E-05 |
| Object vs Person (Dynamic) | 3.60E-02 | 2.00E-08 | 1.70E-04 | 1.80E-03 |
| Object vs Place (Dynamic) | 4.50E-04 | 2.90E-05 | 1.40E-04 | 3.90E-08 |
| Belief vs Photo | 1.00 | 1.00 | 1.00 | 1.00 |
| Sentences vs Nonwords | 1.00 | 1.00 | 1.00 | 3.90E-01 |
| Object vs Person (Visual) | 7.60E-01 | 6.50E-03 | 1.20E-04 | 9.20E-02 |

**Table S5.** *P*-values from object-preferring ROI analysis. Linear mixed model across runs, with participant included as random effect.

|  | **AntTemp** | **PostTemp** | **TPJ** | **IFG** | **MFG** |
| --- | --- | --- | --- | --- | --- |
| Person vs Place (Visual) | 4.70E-01 | 3.50E-01 | 2.90E-05 | 7.30E-02 | 1.50E-03 |
| Person vs Object (Visual) | 5.20E-04 | 8.40E-01 | 2.80E-05 | 1.60E-01 | 2.10E-03 |
| Familiar vs Unfamiliar Face | 1.00E-02 | 8.50E-01 | 5.10E-05 | 1.20E-01 | 2.10E-01 |
| Person vs Place (Semantic) | 1.00 | 1.00 | 1.00 | 1.00 | 4.00E-01 |
| Person vs Object (Semantic) | 2.20E-16 | 2.30E-01 | 2.60E-07 | 9.70E-01 | 1.20E-02 |
| Person vs Place (Episodic) | 3.40E-05 | 3.10E-03 | 2.10E-04 | 1.00E-04 | 6.20E-07 |
| Person vs Object (Episodic) | 3.30E-10 | 3.50E-04 | 2.20E-07 | 1.00E-07 | 4.80E-06 |
| Person vs Place (Dynamic) | 1.00 | 1.30E-02 | 9.50E-04 | 3.00E-05 | 4.20E-02 |
| Person vs Object (Dynamic) | 3.80E-01 | 6.70E-02 | 2.70E-02 | 5.10E-02 | 2.30E-02 |
| Belief vs Photo | 2.20E-01 | 4.80E-01 | 1.10E-04 | 2.00E-01 | 1.00 |
| Sentences vs Nonwords | 6.00E-15 | 2.60E-11 | 6.30E-10 | 1.20E-10 | 7.30E-12 |

**Table S6.** *P*-values from language ROI analysis. Linear mixed model across runs, with participant included as random effect.

|  | **MPC** | **MPFC** | **TPJ** | **STS** | **SFG** | **TP** |
| --- | --- | --- | --- | --- | --- | --- |
| Person vs Place (Visual) | 2.50E-10 | 3.00E-12 | 1.50E-06 | 5.70E-11 | 7.00E-06 | 1.30E-07 |
| Person vs Object (Visual) | 1.50E-11 | 5.30E-19 | 2.90E-09 | 1.50E-11 | 1.90E-12 | 3.10E-10 |
| Familiar vs Unfamiliar Face | 2.50E-07 | 1.40E-11 | 4.00E-06 | 1.80E-08 | 3.10E-07 | 8.70E-08 |
| Person vs Place (Semantic) | 1.40E-11 | 7.50E-11 | 2.00E-07 | 9.90E-09 | 1.90E-07 | 1.10E-13 |
| Person vs Object (Semantic) | 2.10E-11 | 4.10E-13 | 2.40E-10 | 1.70E-09 | 8.10E-11 | 5.30E-15 |
| Person vs Place (Episodic) | 8.80E-08 | 2.70E-06 | 5.40E-03 | 2.10E-05 | 1.70E-02 | 9.90E-09 |
| Person vs Object (Episodic) | 3.60E-09 | 1.80E-10 | 3.90E-08 | 8.90E-16 | 2.70E-05 | 1.30E-09 |
| Person vs Place (Dynamic) | 1.40E-03 | 4.10E-03 | 3.40E-04 | 2.10E-02 | 4.00E-01 | 5.70E-06 |
| Person vs Object (Dynamic) | 1.90E-03 | 2.60E-06 | 1.70E-03 | 1.80E-03 | 3.80E-03 | 1.10E-06 |
| Belief vs Photo | 1.70E-09 | 8.80E-06 | 2.90E-10 | 5.70E-10 | 3.30E-07 | 7.40E-05 |
| Sentences vs Nonwords | 3.90E-01 | 1.00 | 6.40E-01 | 5.00E-06 | 1.00 | 2.60E-03 |

**Table S7.** *P*-values from ToM ROI analysis. Linear mixed model across runs, with participant included as random effect.

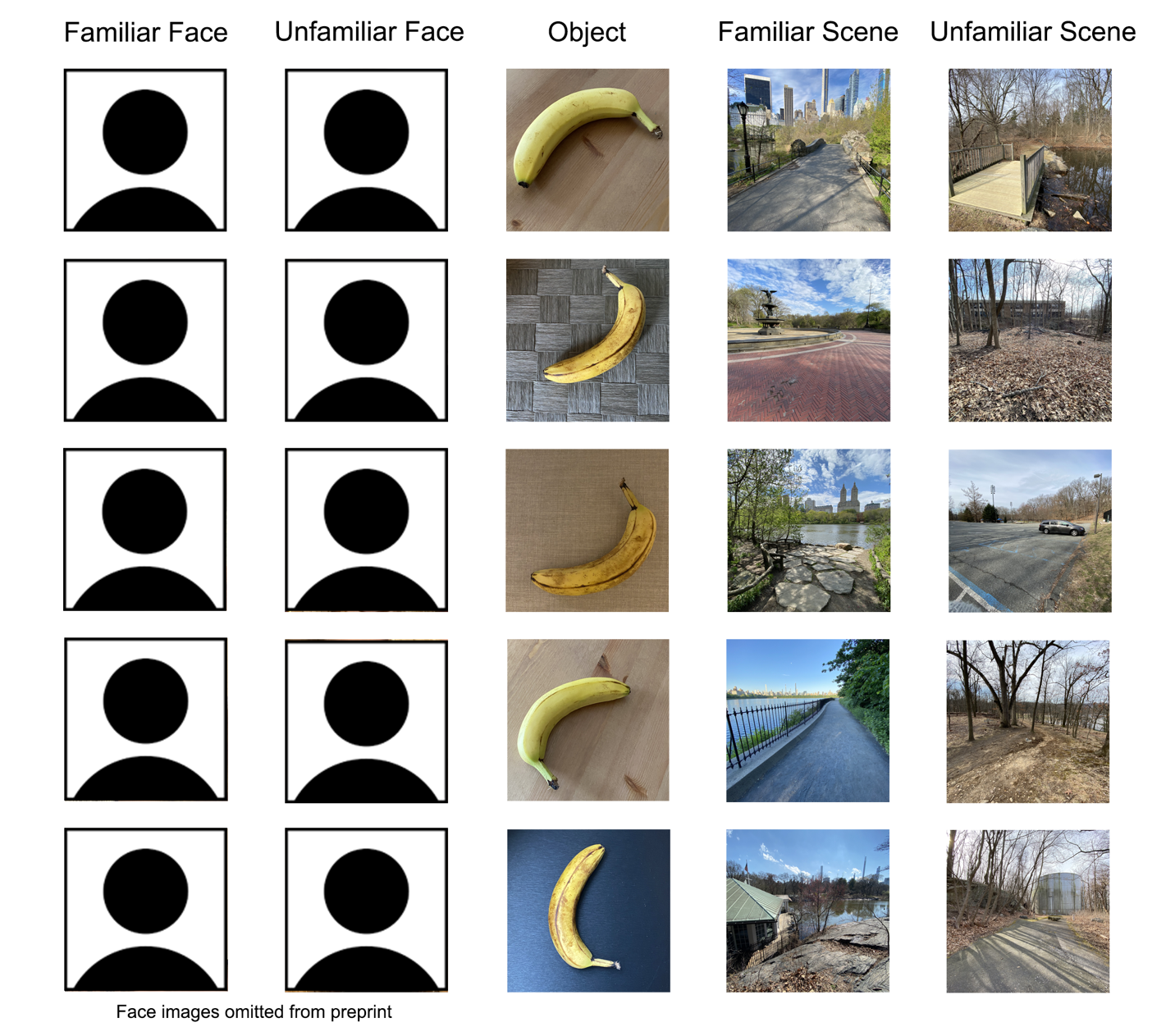

**Figure S1.** Additional examples of images used in the visual perception task. Each row contains examples from a given participant, condition, and identity.

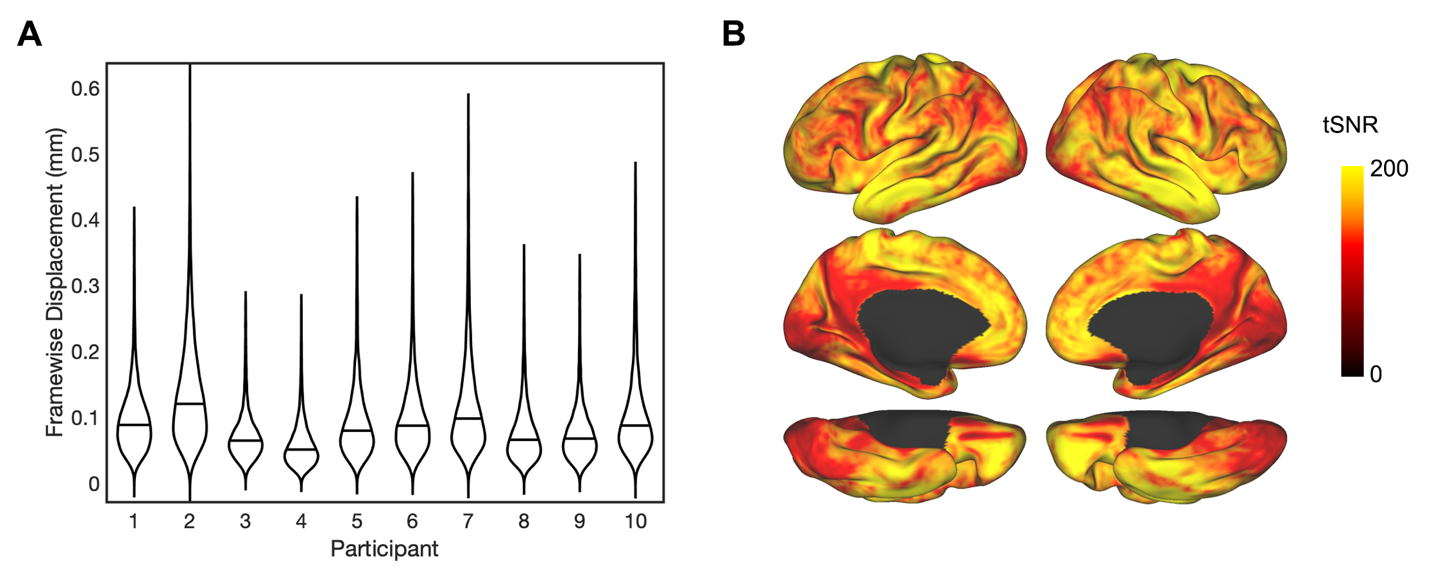

**Figure S2.** Data quality metrics. **A)** Violin plots of framewise displacement distributions for each participant, with the median value marked. For visualization purposes, we removed outliers defined as values greater than the upper quartile plus five times the interquartile range (.6% of values). **B)** Temporal signal-to-noise ratio (tSNR) of task fMRI data, averaged across participants and datasets.

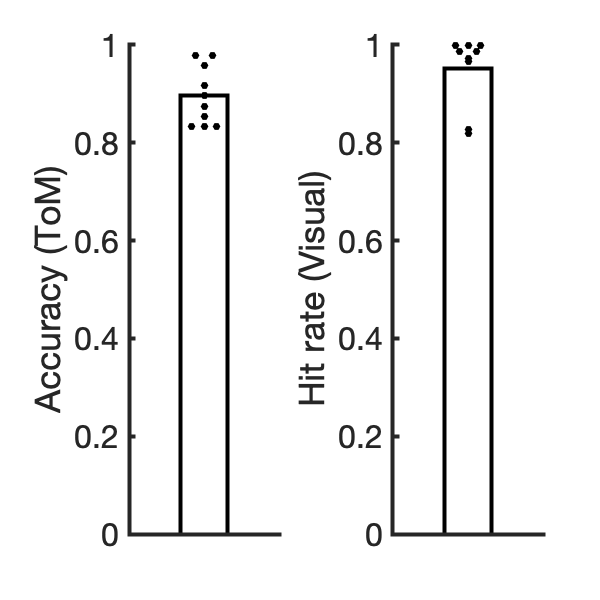

**Figure S3.** Scanner behavioral performance. Left: Accuracy on true/false questions in the theory of mind (ToM) experiment. Right: Hit rate on the one-back task in the visual experiment. Dots show performance of individual participants.

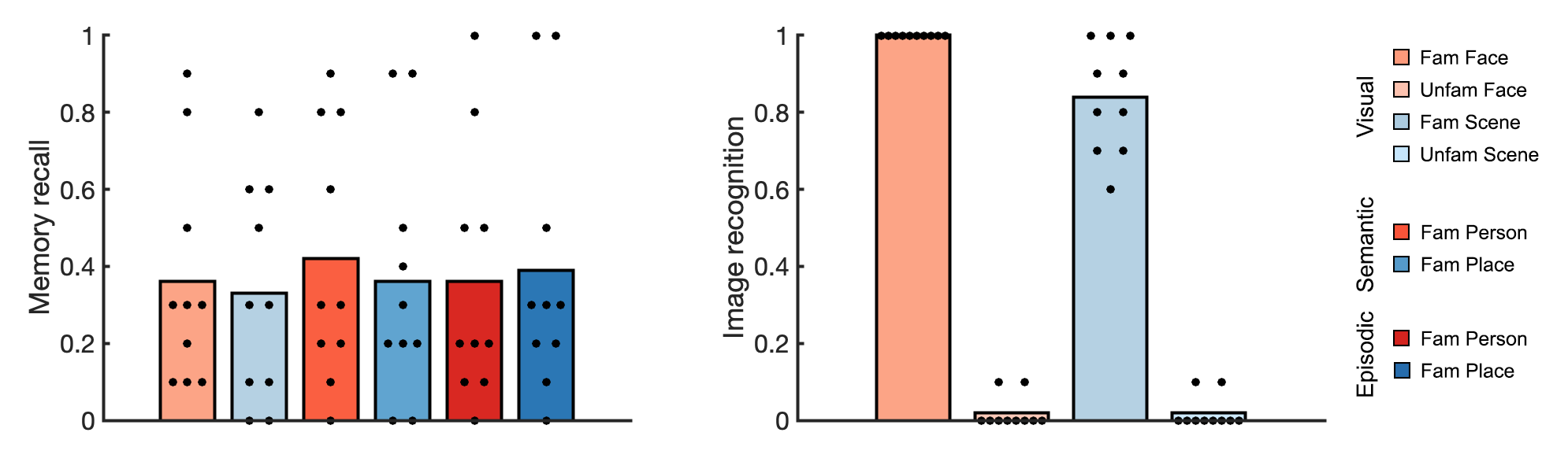

**Figure S4.** Post-scan questionnaire responses. Left: Participants were asked what proportion of time they spent recalling memories of past events during the familiar person and place conditions of visual, semantic, and episodic tasks. Right: Participants were asked what proportion of images they recognized in the familiar and unfamiliar person and place conditions of the visual task. Dots show responses in individual participants. In the small number of cases where participants reported recognizing more than 0% of unfamiliar images, they were probed further, and revealed that they did not in fact recognize the particular person or place displayed, but thought the images looked familiar.

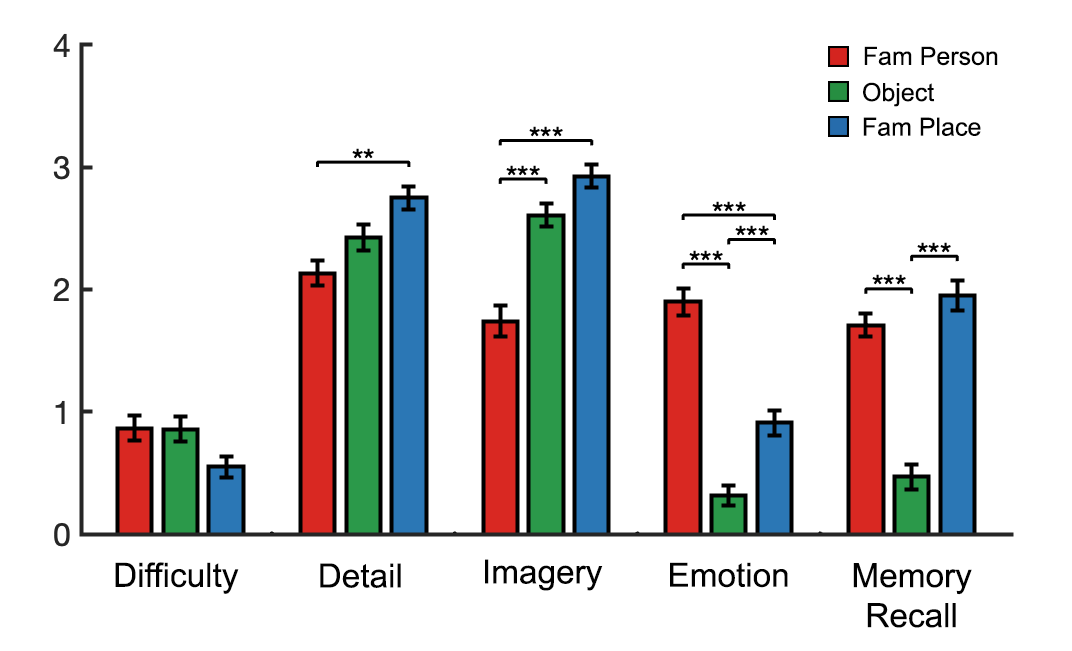

**Figure S5.** Post-scan behavioral ratings of imagined events in the episodic task, including ratings of difficulty, level of detail, visual imagery, experience of emotion, and memory recall. Error bars show standard error across items. ** *P* < 10^-3^, *** *P* < 10^-4^ (linear mixed model across items, with random intercepts for participant).

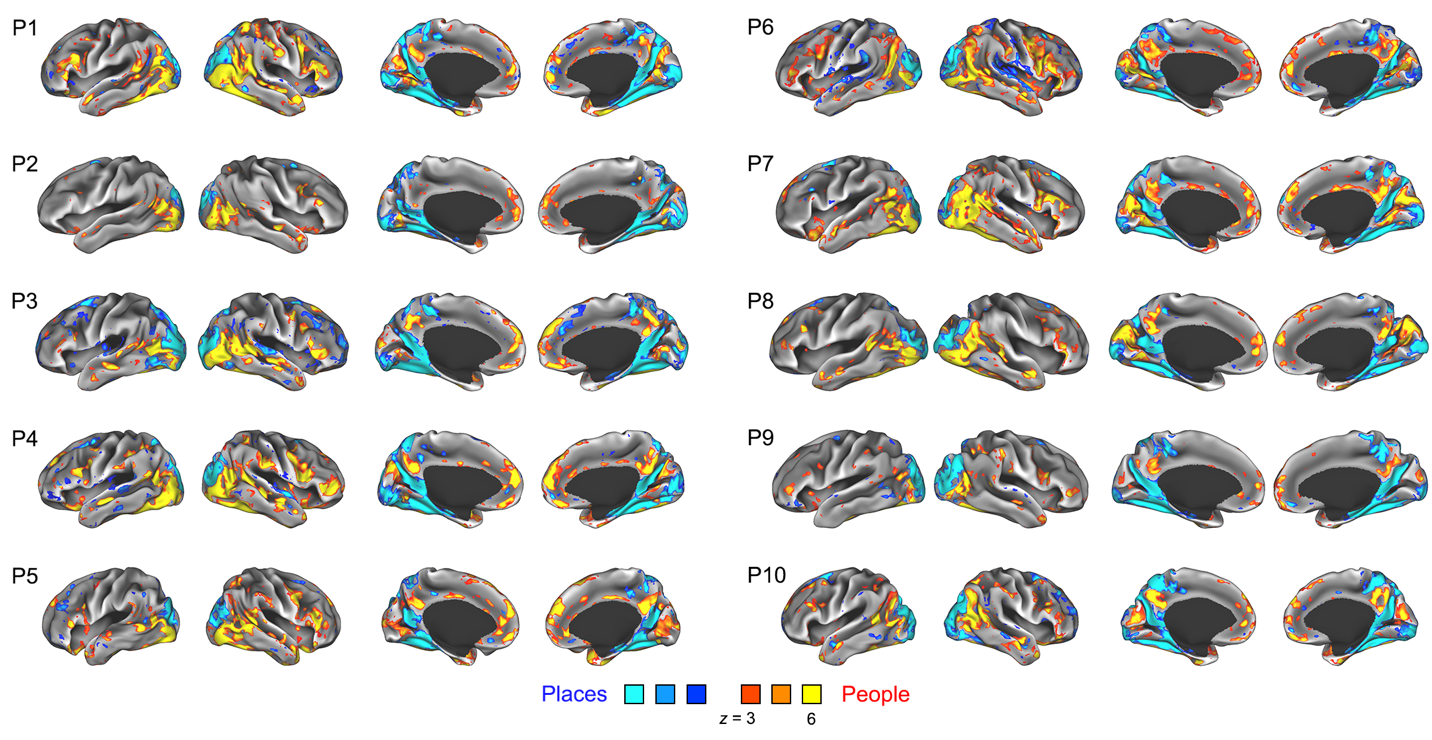

**Figure S6.** Whole-brain general linear model-based responses to people versus places (visual task), across all participants. Results thresholded at a False Discovery Rate of *q* < .01 to correct for multiple comparisons across coordinates.

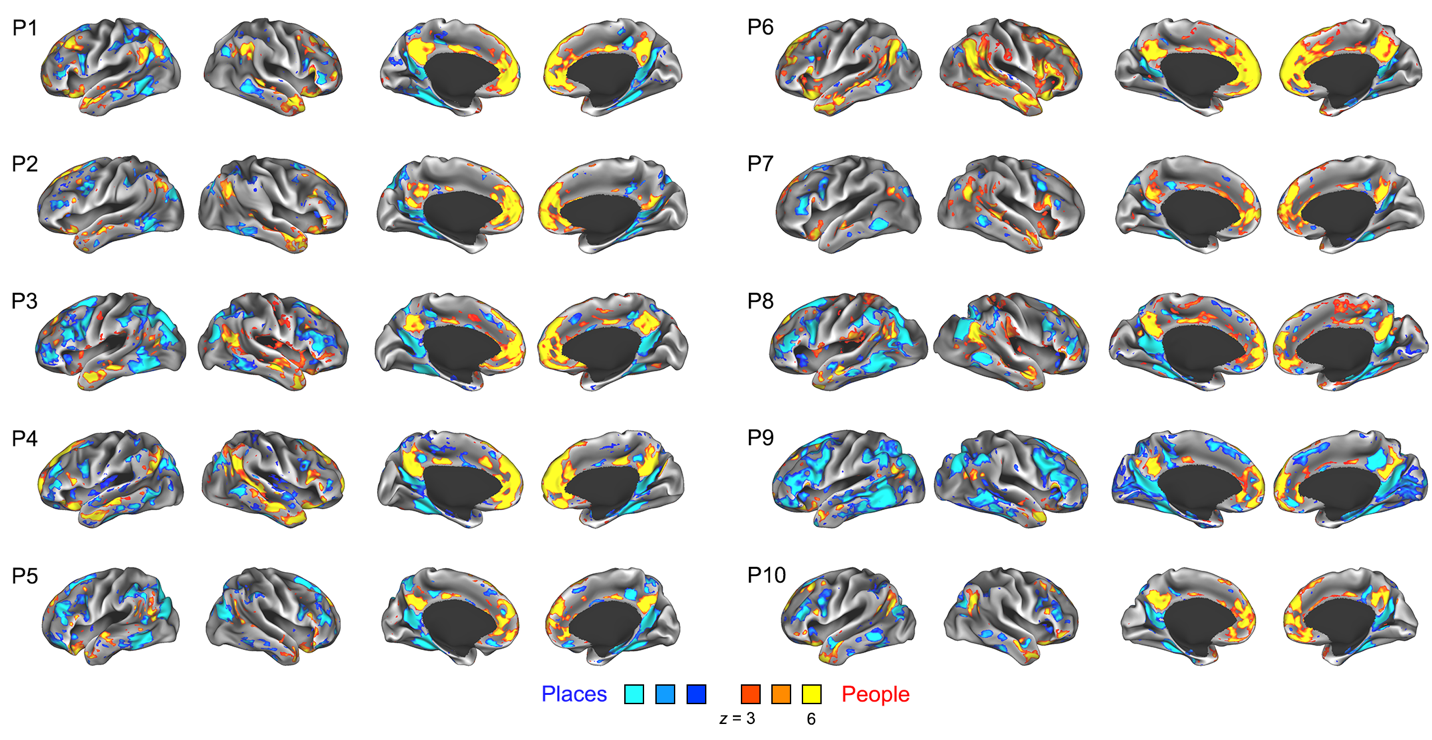

**Figure S7.** Whole-brain general linear model-based responses to people versus places (semantic task), across all participants. Results thresholded at a False Discovery Rate of *q* < .01 to correct for multiple comparisons across coordinates.

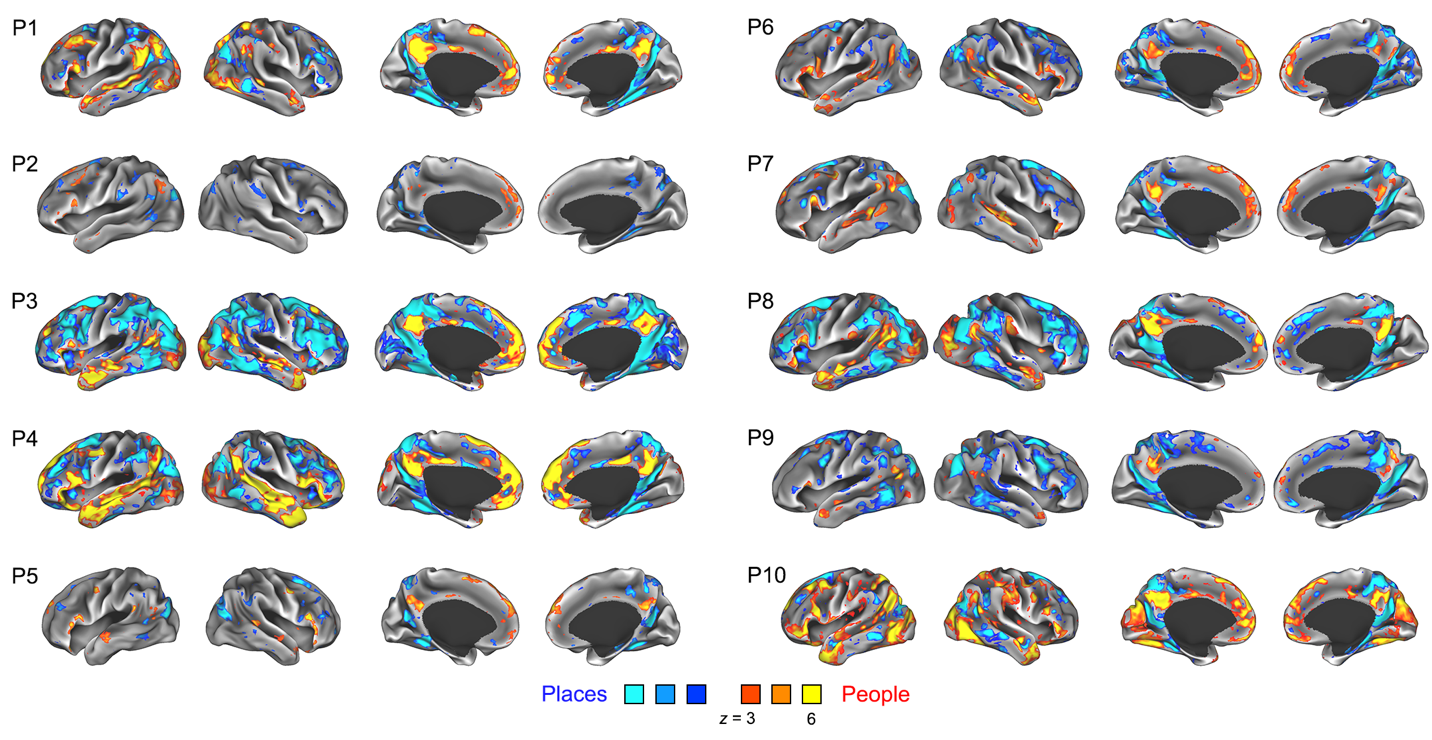

**Figure S8.** Whole-brain general linear model-based responses to people versus places (episodic task), across all participants. Results thresholded at a False Discovery Rate of *q* < .01 to correct for multiple comparisons across coordinates.

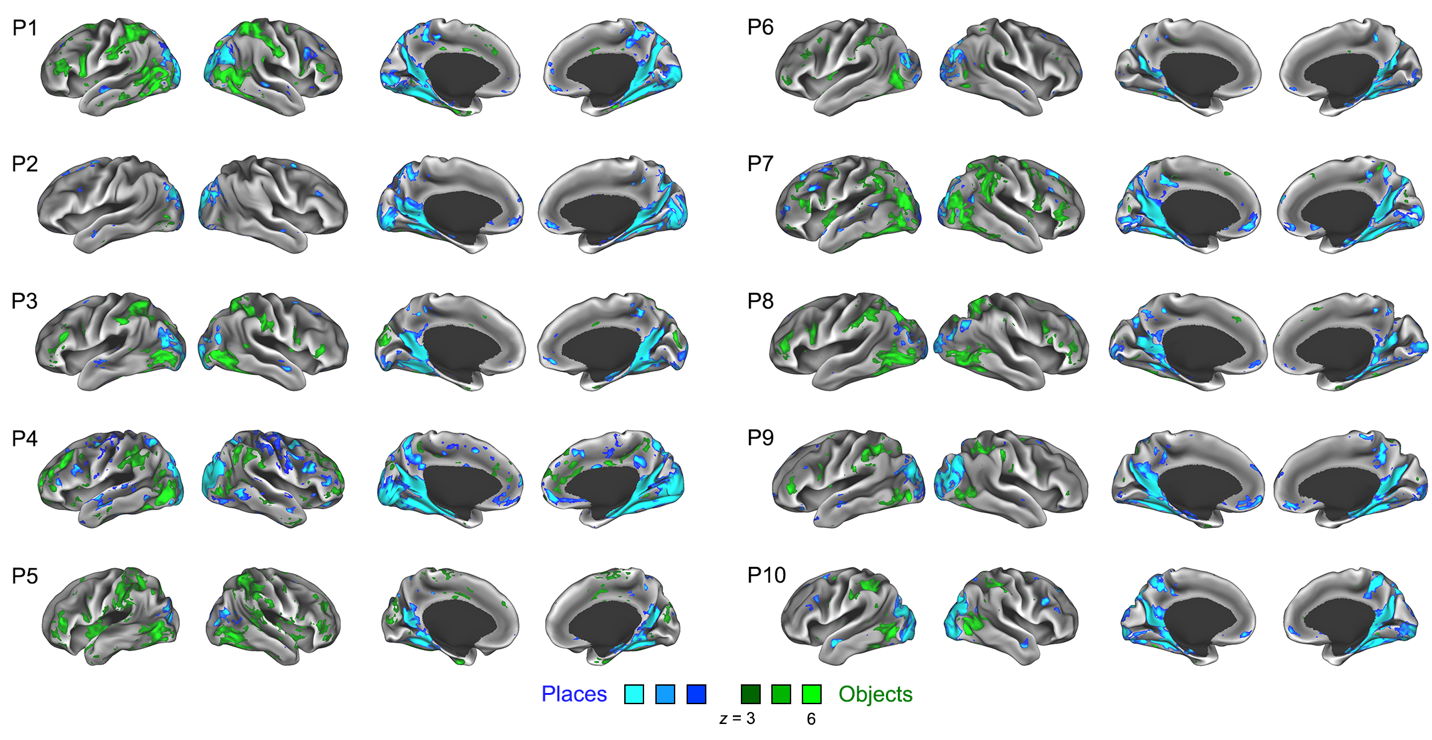

**Figure S9.** Whole-brain general linear model-based responses to objects versus places (visual task), across all participants. Results thresholded at a False Discovery Rate of *q* < .01 to correct for multiple comparisons across coordinates.

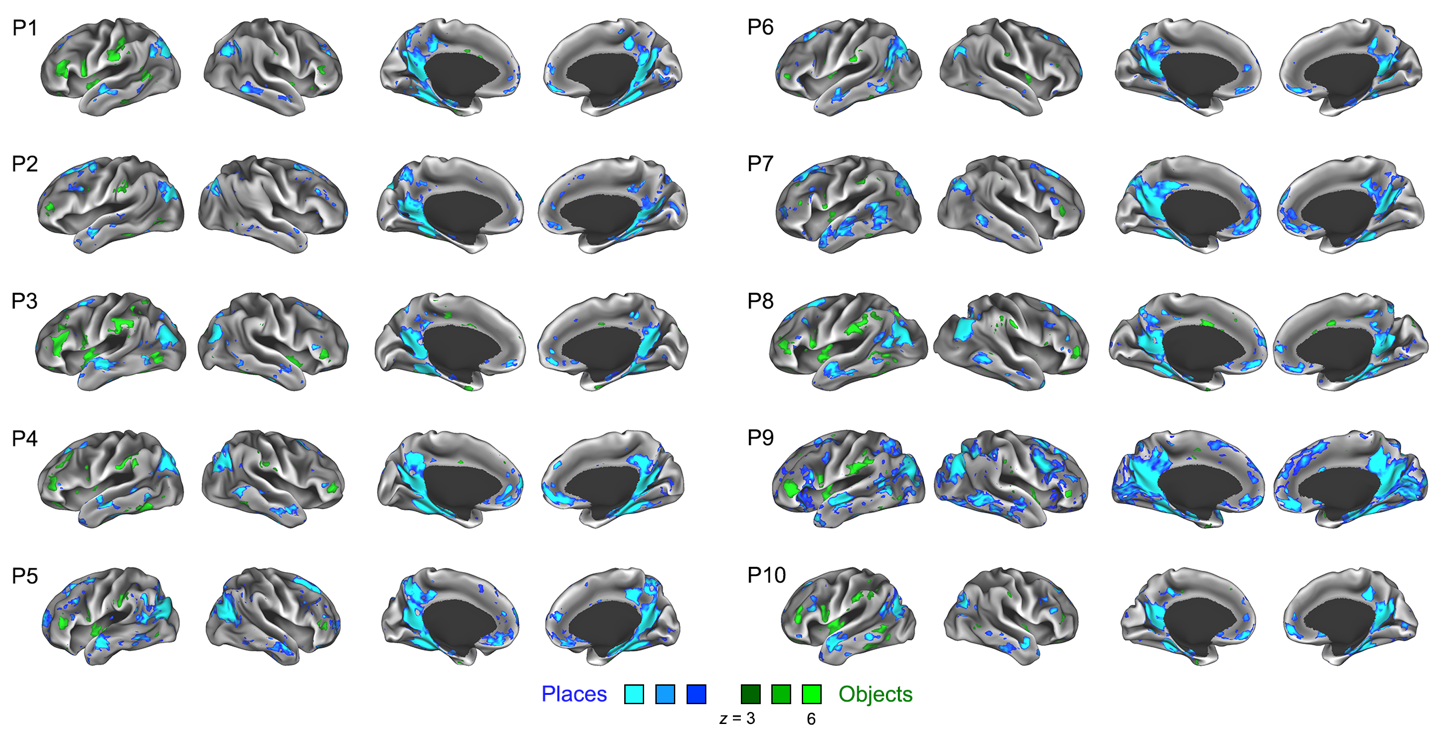

**Figure S10.** Whole-brain general linear model-based responses to objects versus places (semantic task), across all participants. Results thresholded at a False Discovery Rate of *q* < .01 to correct for multiple comparisons across coordinates.

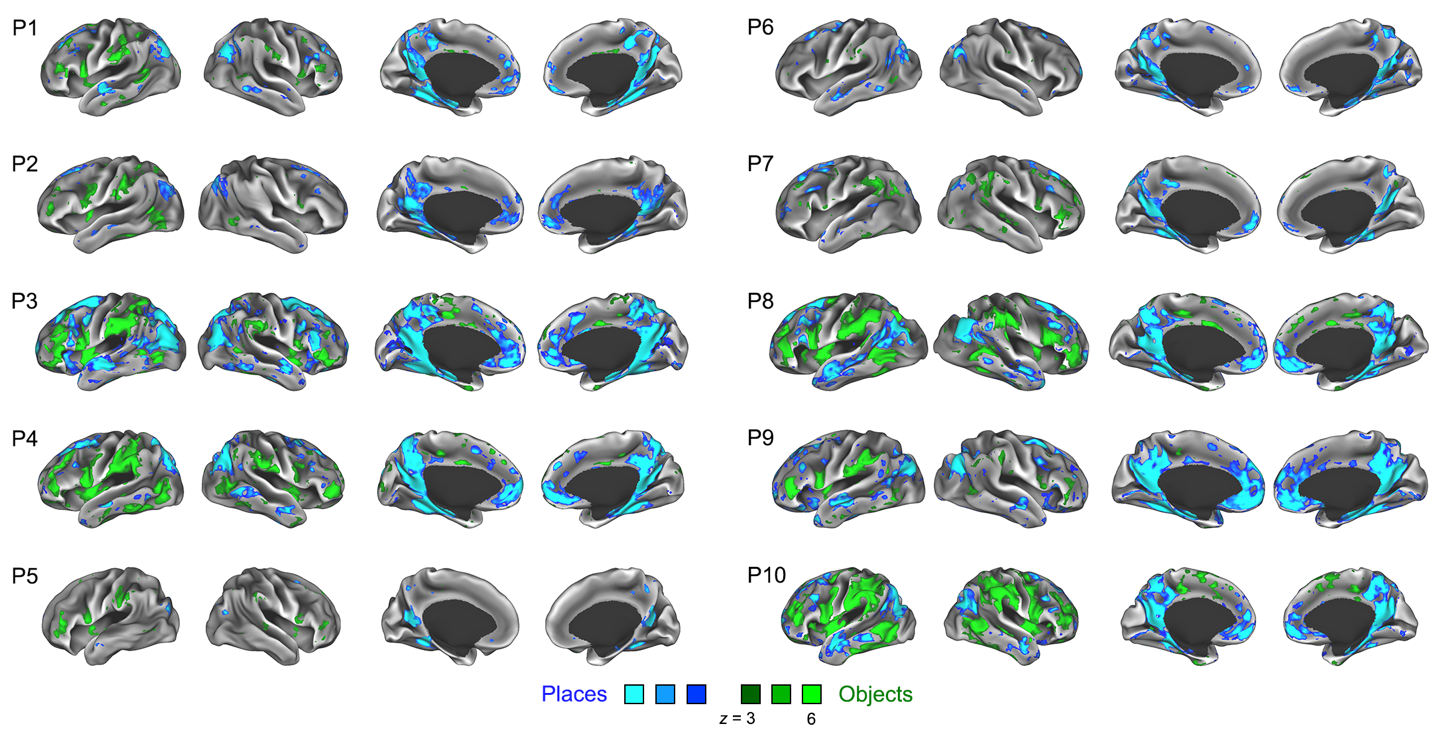

**Figure S11.** Whole-brain general linear model-based responses to objects versus places (episodic task), across all participants. Results thresholded at a False Discovery Rate of *q* < .01 to correct for multiple comparisons across coordinates.

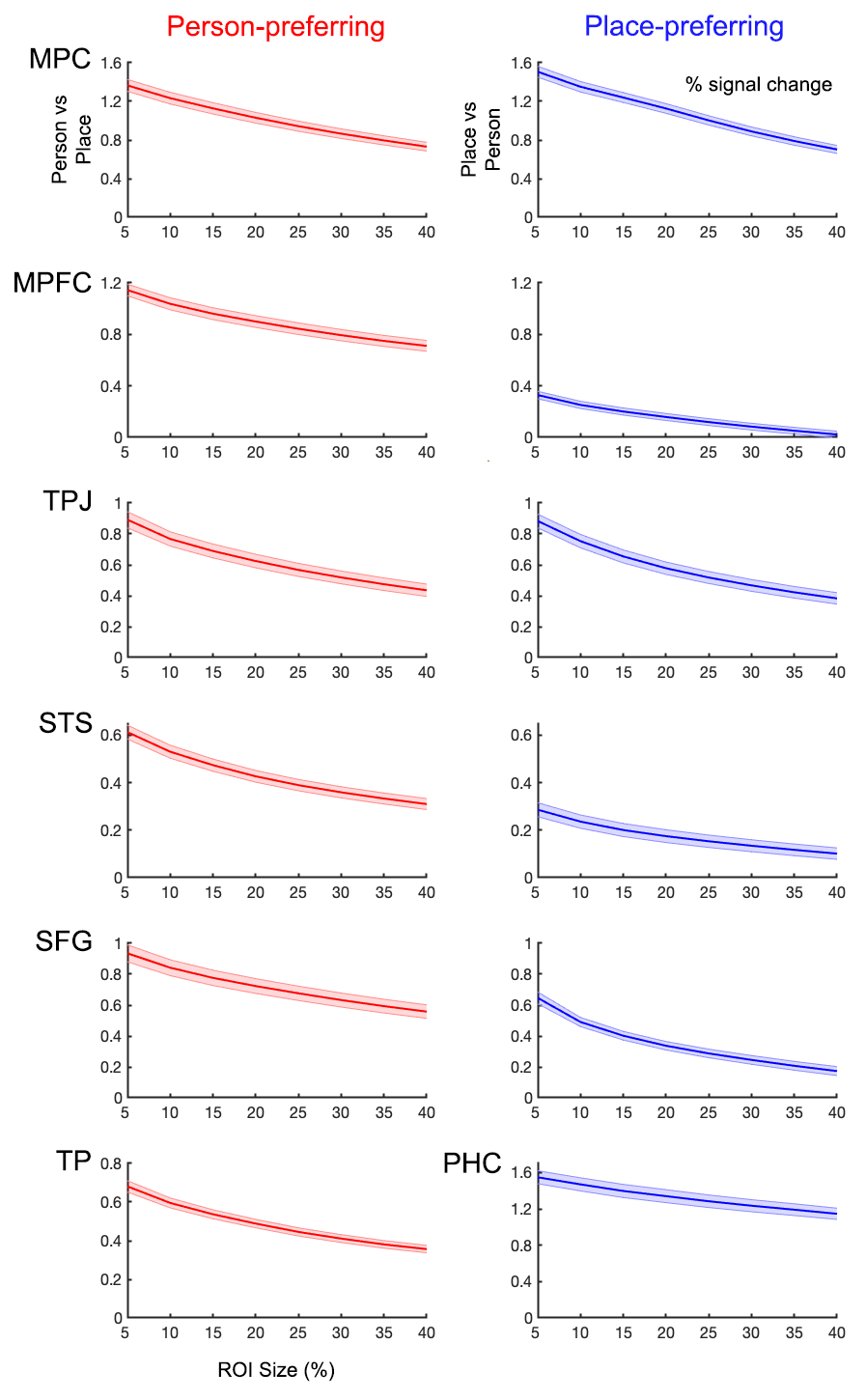

**Figure S12:** Region-of-interest (ROI) responses, across a range of ROI sizes. ROIs were defined as the top N% of person- or place-preferring coordinates (semantic task) within anatomical search spaces, with N varying from 5 to 40%. The response to people over places (or places over people) was extracted in independent data from the same task, and plotted as a function of ROI size. Shaded regions show standard error across runs.

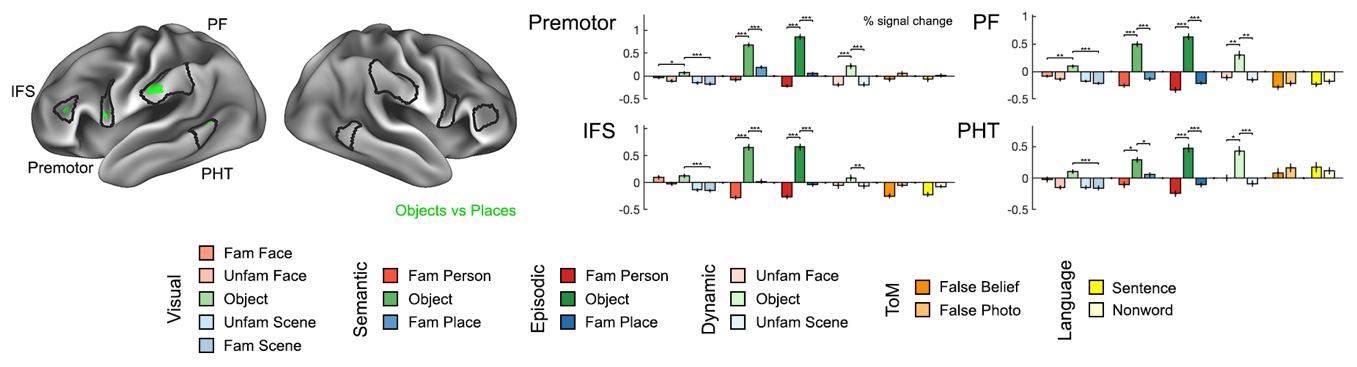

**Figure S13:** Object-sensitive ROI analysis. Left: search spaces and functional ROIs from one representative participant. ROIs were defined as the top 5% of object-sensitive coordinates (object versus place contrast) within each search space. Right: Responses (% signal change) extracted from functionally defined ROIs. Error bars show standard error across runs. * *P* < .05/7 = .0071, ** *P* < 10^-3^, *** *P* < 10^-4^ (linear mixed model across runs, with participant included as random effect). Abbreviations: IFS, interior frontal sulcus; PF, area PF; PHT, area PHT.

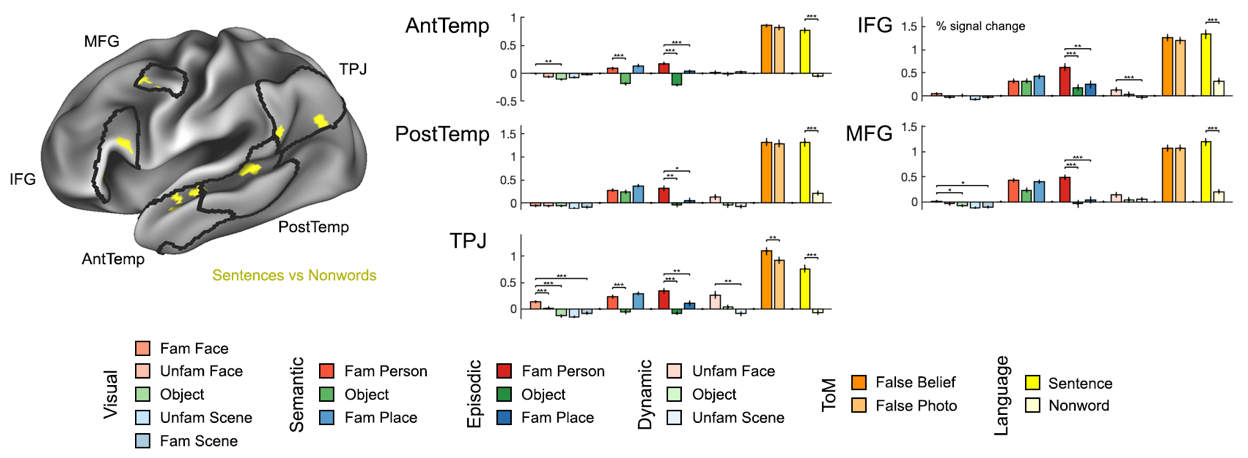

**Figure S14:** Language ROI analysis. Left: search spaces and functional ROIs from one representative participant (for language ROIs, search spaces were restricted to the left hemisphere). ROIs were defined as the top 5% of language-sensitive coordinates (sentences vs nonwords contrast) within each search space. Right: Responses (% signal change) extracted from functionally defined ROIs. Error bars show standard error across runs. * *P* < .05/7 = .0071, ** *P* < 10^-3^, *** *P* < 10^-4^ (linear mixed model across runs, with participant included as random effect). Abbreviations: AntTemp, anterior temporal; IFG, inferior frontal gyrus; MFG, middle frontal gyrus; PostTemp, posterior temporal; TPJ, temporo-parietal junction.

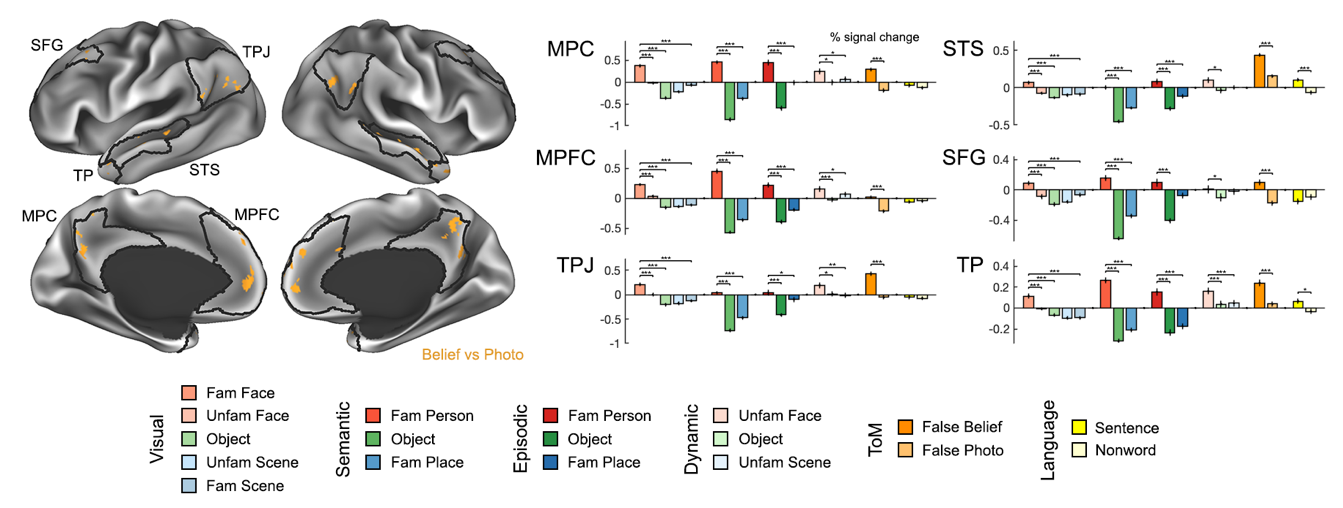

**Figure S15:** Theory of mind ROI analysis. Left: search spaces and functional ROIs from one representative participant. ROIs were defined as the top 5% of ToM-sensitive coordinates (false belief vs false photo contrast) within each search space. Right: Responses (% signal change) extracted from functionally defined ROIs. Error bars show standard error across runs. * *P* < .05/7 = .0071, ** *P* < 10^-3^, *** *P* < 10^-4^ (linear mixed model across runs, with participant included as random effect). Abbreviations: MPC, medial parietal cortex; MPFC, medial prefrontal cortex; SFG, superior frontal gyrus; STS, superior temporal sulcus; TPJ, temporo-parietal junction; TP, temporal pole.

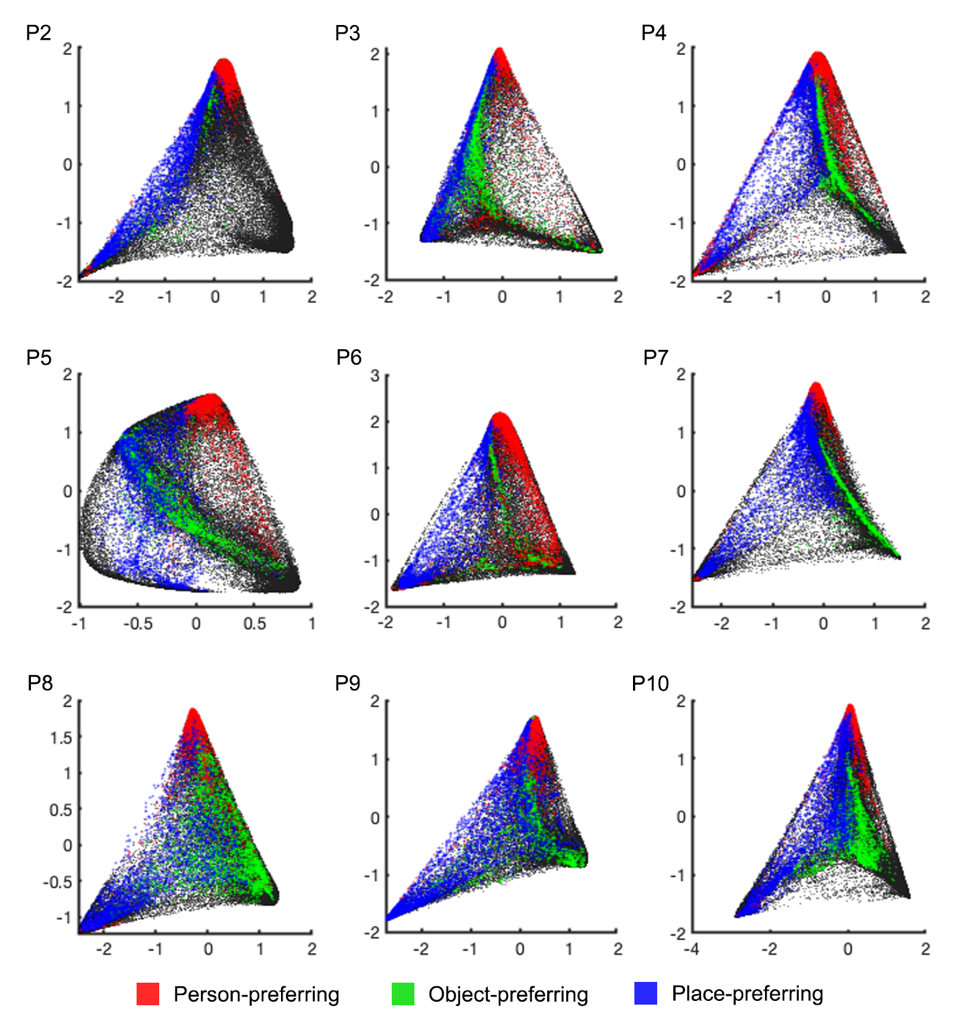

**Figure S16.** Person-, place-, and object-preferring regions (coordinate-wise *Z* > 2.3 across visual, semantic, and episodic tasks), overlaid on functional connectivity distance space, for participants 2-10. The space was defined as a diffusion embedding of individual participant resting-state functional connectivity distances, transformed to the first participant’s space using a Procrustes transform.

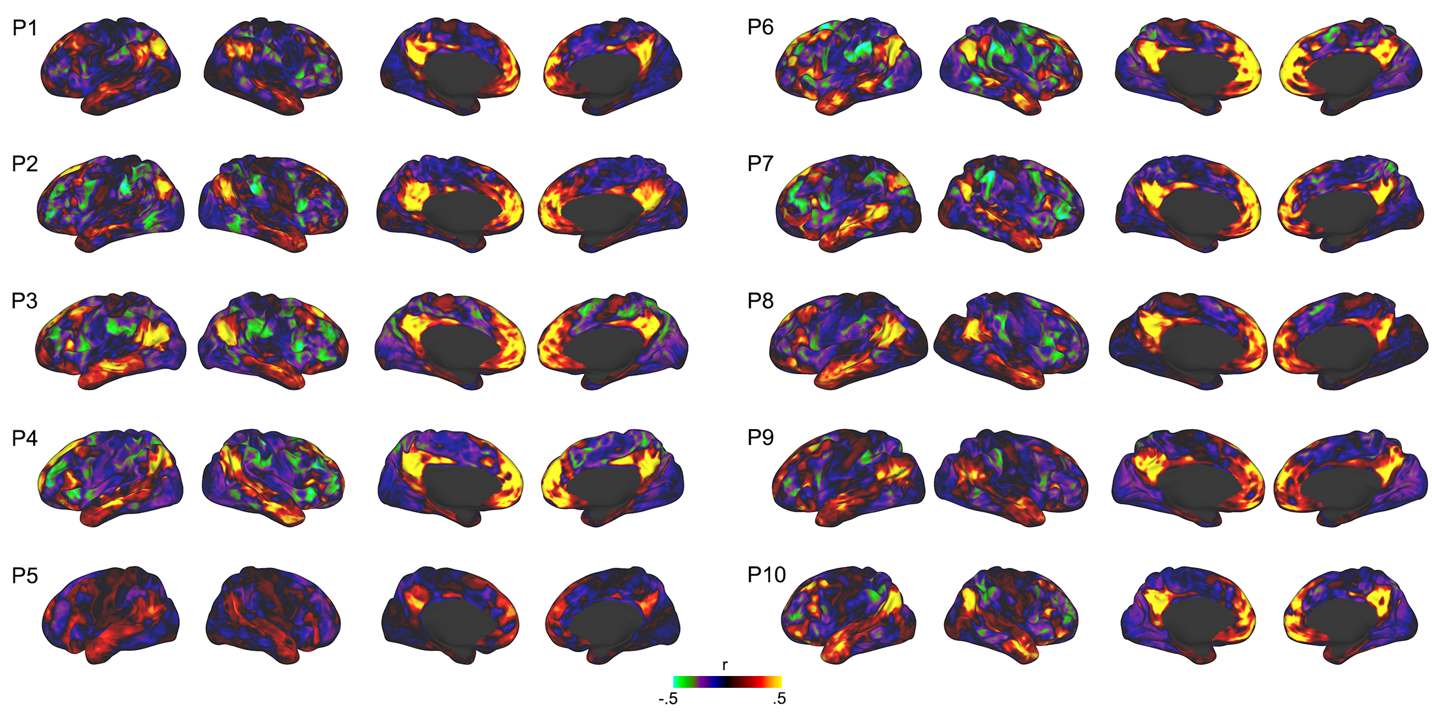

**Figure S17.** Whole-brain resting-state correlation maps from a person-preferring LMPC seed region, across all participants.

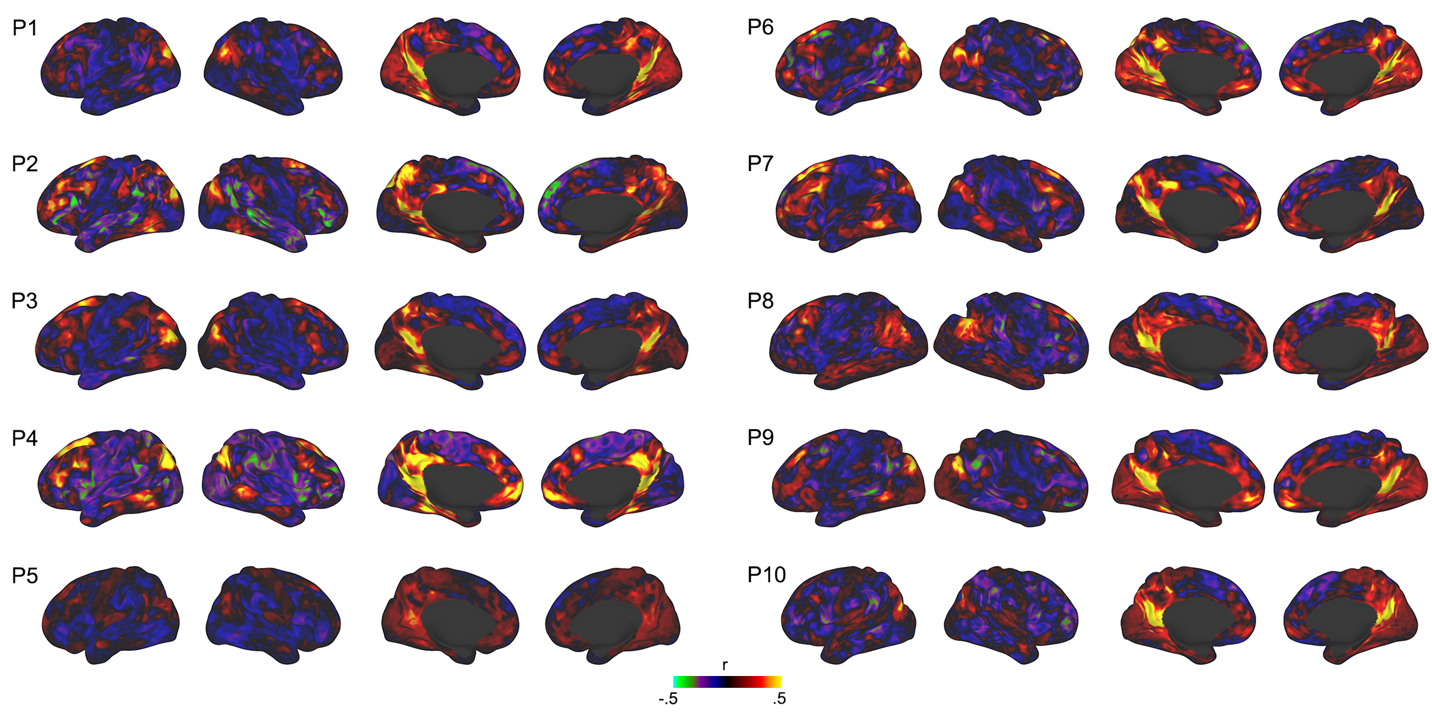

**Figure S18.** Whole-brain resting-state correlation maps from a place-preferring LMPC seed region, across all participants.

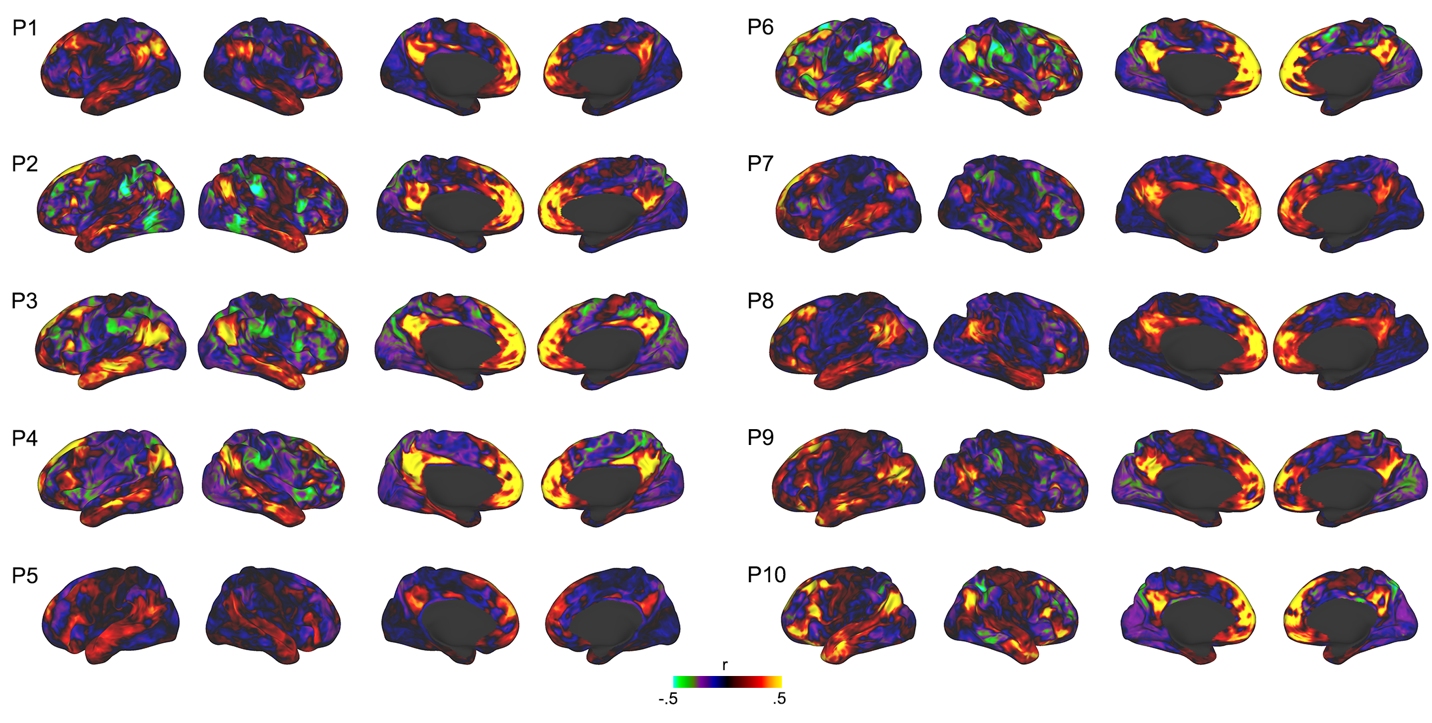

**Figure S19.** Whole-brain resting-state correlation maps from a person-preferring LSFG seed region, across all participants.

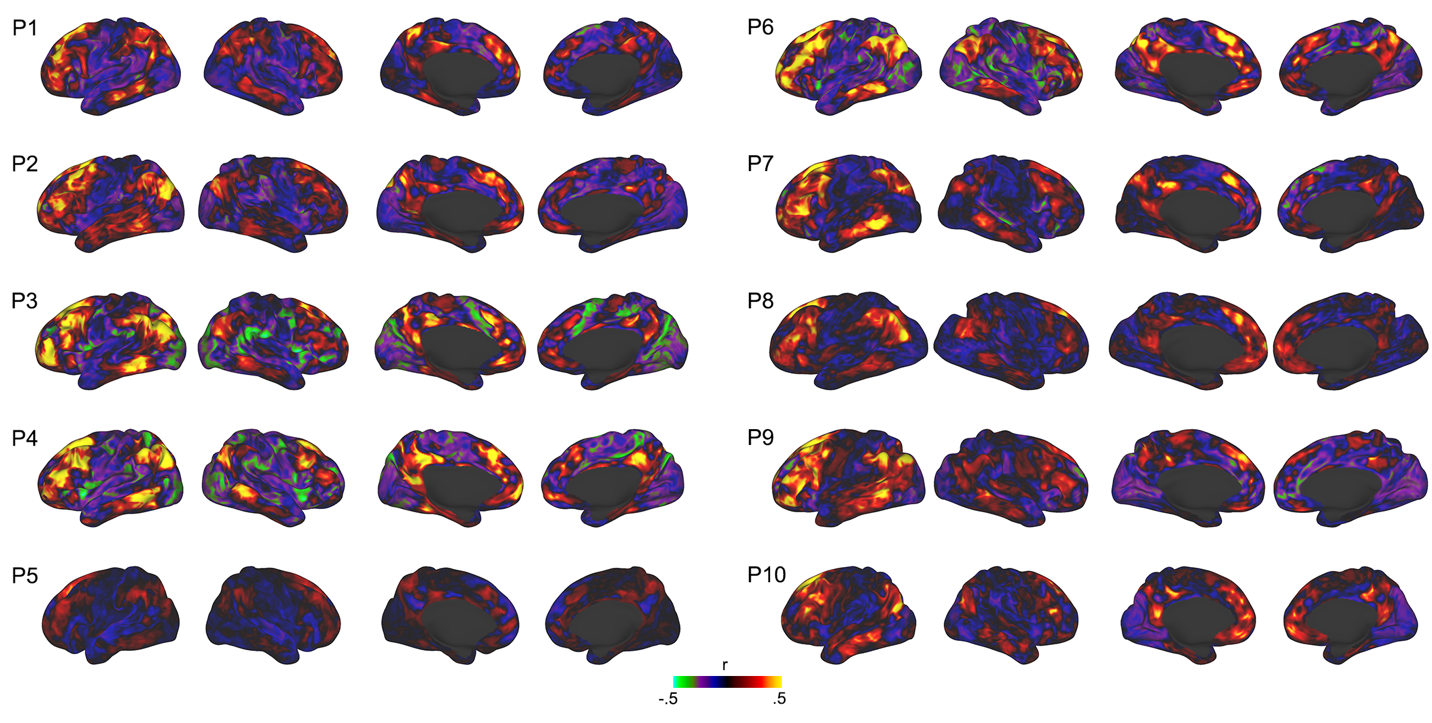

**Figure S20.** Whole-brain resting-state correlation maps from a place-preferring LSFG seed region, across all participants.
